# State-Transition Analysis of Time-Sequential Gene Expression Identifies Critical Points That Predict Leukemia Development

**DOI:** 10.1101/238923

**Authors:** Russell C. Rockne, Sergio Branciamore, Jing Qi, David Frankhouser, Denis O’Meally, Wei-Kai Hua, Guerry J. Cook, Emily Carnahan, Lianjun Zhang, Ayelet Marom, Herman Wu, Davide Maestrini, Xiwei Wu, Yate-Ching Yuan, Zheng Liu, Leo D. Wang, Stephen J. Forman, Nadia Carlesso, Ya-Huei Kuo, Guido Marcucci

## Abstract

Temporal dynamics of gene expression are informative of changes associated with disease development and evolution. Given the complexity of high-dimensional temporal datasets, an analytical framework guided by a robust theory is needed to interpret time-sequential changes and to predict system dynamics. Herein, we use acute myeloid leukemia as a proof-of-principle to model gene expression dynamics in a transcriptome state-space constructed based on time-sequential RNA-sequencing data. We describe the construction of a state-transition model to identify state-transition critical points which accurately predicts leukemia development. We show an analytical approach based on state-transition critical points identified step-wise transcriptomic perturbations driving leukemia progression. Furthermore, the gene(s) trajectory and geometry of the transcriptome state-space provides biologically-relevant gene expression signals that are not synchronized in time, and allows quantification of gene(s) contribution to leukemia development. Therefore, our state-transition model can synthesize information, identify critical points to guide interpretation of transcriptome trajectories and predict disease development.

**Graphical Abstract:** 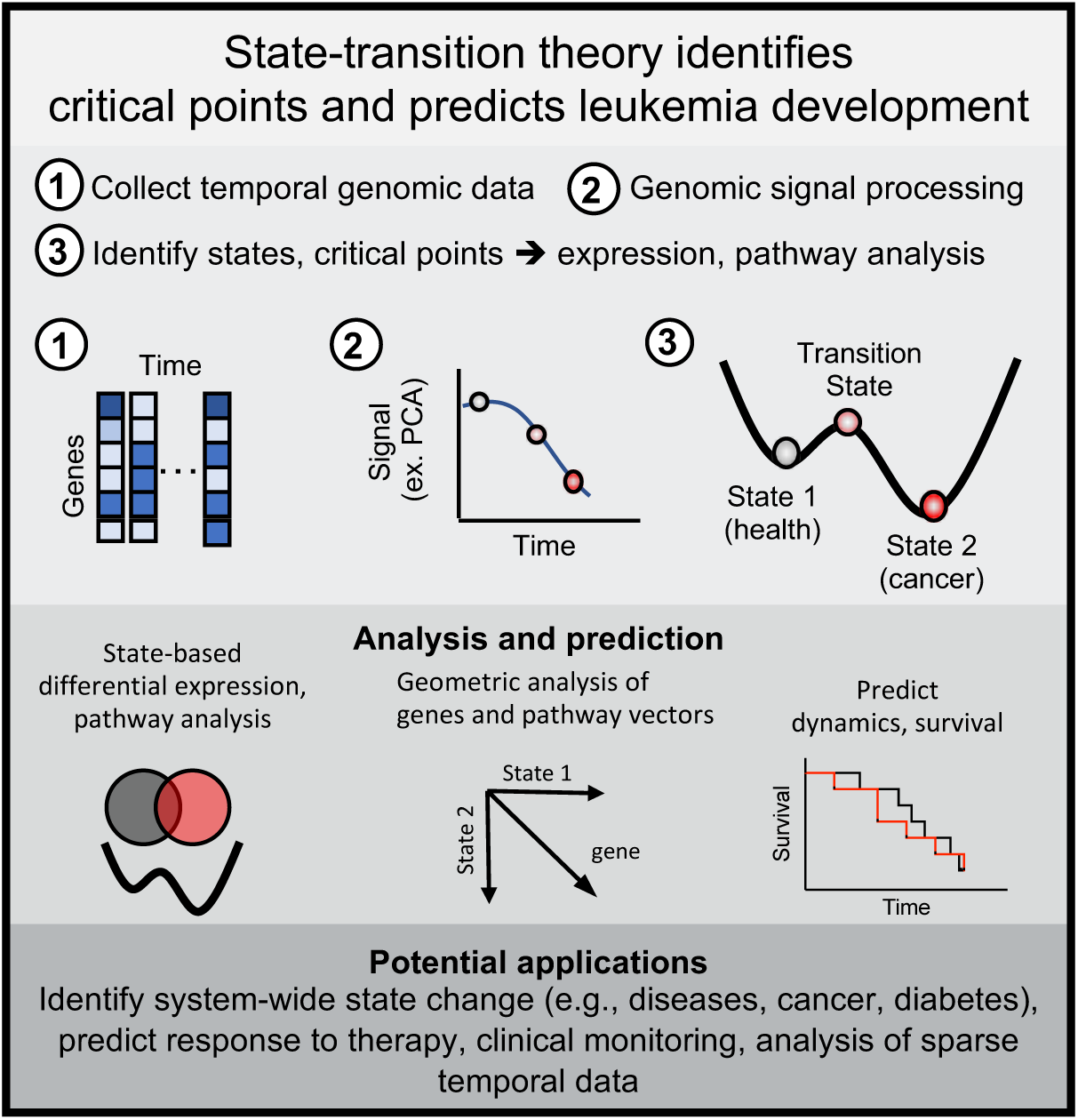

**In brief:** The theory of state-transition is applied to acute myeloid leukemia (AML) to model transcriptome dynamics and trajectories in a state-space, and is used to identify critical points corresponding to critical transcriptomic perturbations that predict leukemia development.

**Highlights:**

- Leukemia transcriptome dynamics are modeled as movement in transcriptome state-space
- State-transition model and critical points accurately predicts leukemia development
- Critical point-based approach identifies step-wise transcriptome events in leukemia
- State-based geometric analysis provides quantification of leukemogenic contribution

## Introduction

A complex disease such as cancer evolves as a dynamic system wherein multilayer interconnected inputs collectively produce a disease state corresponding to a clinical phenotype. Identification of genomic alterations including gene mutations, epigenetic changes and gene expression profiles by high-throughput sequencing assays are becoming a part of the routine clinical assessment of cancer patients at diagnosis and subsequent follow-ups. With the potential for tens of thousands of statistically significant genomic alterations to be detected at any given time point, it is challenging to quantitatively determine which changes are biologically relevant. In addition, these profiles continuously change over time as result of malignant cell transformation, cancer evolution, and ultimately, response to treatment. The analysis and interpretation of such a large volume of data comprising dynamic changes with regard to biological and clinical evolution poses a significant challenge. Although methods are available to analyze time-series genomic data(Bar-Joseph et al., 2012; Sanavia et al., 2015; Spies et al., 2017), to date it has been difficult to meaningfully interpret global gene expression changes over time and use them to predict system dynamics. To achieve the ultimate goal of predicting cancer system dynamics, it is of critical biological and clinical importance to develop a framework guided by a robust theory by which to analyze temporal genomic data.

A central challenge for interpreting time-sequential changes in gene expression is the identification and prioritization of the genes, pathways, and sampling timepoints that are necessary for understanding changes of interest (e.g., transition from health to cancer, cancer evolution, response to treatments). In theory, if we could continuously monitor the state of health of an individual through longitudinal data collection, we would observe critical inflection points at which the system transitions from one state to another (e.g., from health to cancer, from localized to metastatic disease, from therapy responsiveness to therapy refractoriness). In such a hypothetical scenario, any number of existing methods would suffice to analyze and identify specific moments in time—and corresponding genomic, clonal, or immunological events—critical to cancer dynamics. In practice, however, data are typically sparsely collected over time, and a system is unlikely to be observed precisely at the relevant critical points. Thus, existing genomic analysis approaches are limited in their ability to identify relevant alterations and interpret temporal dynamics in many real-world situations.

Here, we present an analytical approach based on state-transition theory, which allows us to conceptually fill in the gaps of incomplete and temporally sparse genomic data and identify time-dependent critical transcriptomic perturbations that predict cancer progression. We illustrate how state-transition theory can be applied to model and predict the development of acute myeloid leukemia (AML) based on time-series RNA-sequencing data collected at sparse timepoints. AML is a devastating malignancy of the hematopoietic system that may rapidly lead to bone marrow failure and death. Approximately 21,000 new patients are diagnosed with AML each year in the United States, and the latest 5-year overall survival rate remains at only 28% (https://seer.cancer.gov)(Noone et al., 2018). AML comprises multiple entities characterized by gene mutations and chromosomal abnormalities that drive leukemogenesis and predict prognosis and/or therapeutic response.^11^ Genomic studies such as the cancer genome atlas have revealed various mutational landscapes in AML, highlighting patterns of cooperation and exclusivity among the gene mutations(Dohner et al., 2015). These gene mutations ultimately alter the expression of downstream target genes and are therefore associated with unique gene expression profiles representing functional networks in leukemic cell biology. Thus, changes of the global system-level gene expression (i.e., the transcriptome) may be viewed as an *emergent property of the system*, which is informative with regard to disease evolution.

To model AML development as a state-transition, we use a well-established conditional knock-in mouse model that mimics a subset of human AML driven by the fusion gene *CBFB-MYH11* (CM), corresponding to the cytogenetic rearrangement inv(16)(p13.1q22) or t(16;16)(p13.1;q22) [henceforth inv(16)]. Inv(16) is one of the most common recurrent cytogenetic aberrations and is found in approximately 5-12% of all patients with AML. Induction of CM expression disrupts normal hematopoietic differentiation, resulting in perturbed hematopoiesis in the bone marrow and an increased probability of state-transition from health to leukemia. We show here that temporal transcriptome dynamics in peripheral blood mononuclear cells (PBMCs) from CM mice are predictive of state progression represented as the movement of a transcriptome-particle in a two-dimensional state-space (i.e., normal hematopoiesis and leukemia). Critical points of these transcriptome trajectories in the state-space correspond to specific gene expression changes that contribute to the progression from a reference state of normal hematopoiesis to leukemia, thereby providing biological insights into critical time-dependent transcriptomic pertubations during leukemogenesis.

## Results

### State-transition dynamics of the transcriptome

State-transition theory has a rich mathematical foundation(Pavliotis, 2014) and has been broadly applied in various scientific fields (i.e., chemistry, physics, and biology(Esteban et al., 2018; Folguera-Blasco et al., 2018; Herring et al., 2017; Hormoz et al., 2016; Pastushenko et al., 2018; Zhou et al., 2012)). To our knowledge, it has not yet been applied to study cancer progression from a reference state of health (i.e., normal hematopoiesis) to a state of disease (i.e., leukemia). To apply state-transition theory to this setting, we started from the observation that the composition of PBMCs changes at different stages of leukemia development, treatment response, or post-treatment disease relapse. We considered blood rather than bone marrow as the organ of interest for our study because it is much more accessible for frequent sequential sampling and therefore our approach, if successful, may be easily applied in the clinical setting.

Changes in the cellular composition of PBMCs are obvious once disease is clinically present (defined here as at least 20% circulating blasts). However, current approaches (including diagnostic molecular testing) may not fully detect the subtle changes occurring in between time-sequential sampling before overt leukemia is present, and therefore unable to predict with accuracy the trajectory of the disease development or regression at any discrete timepoint. Herein, we hypothesize that the PBMC transcriptome can be modeled by representing it as a particle undergoing Brownian motion in a double-well quasi-potential that determines the probability of the system to transition from a state of health to a state of leukemia (Figure 1A). To represent health to leukemia transition, we postulated that in a normal hematopoiesis state, a large energy barrier reduces the probability of the system to transition to a leukemia state. However, once hematopoiesis is perturbed by the expression of a leukemogenic event (i.e., the fusion gene *CM* expression in the case of our murine AML model), the double-well quasi-potential energy landscape is altered in a way that the energy barrier is lowered and the probability of transition from normal hematopoiesis to leukemia significantly increases (Figure 1B). Mathematically, this double-well quasi-potential can be represented as a 4^th^ degree polynomial 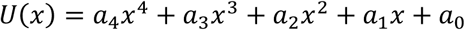 with constant coefficients *α_i_*. The equation of motion of the particle in the double-well quasi-potential then takes the form of a stochastic differential equation 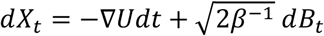 where *X_t_* denotes the location of the particle at time *t* and *dB_t_* is a Brownian motion that is uncorrelated in time 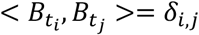, with *δ_i,j_* being the Dirac delta function and *β*^−1^ is the diffusion coefficient.

**Figure 1.**
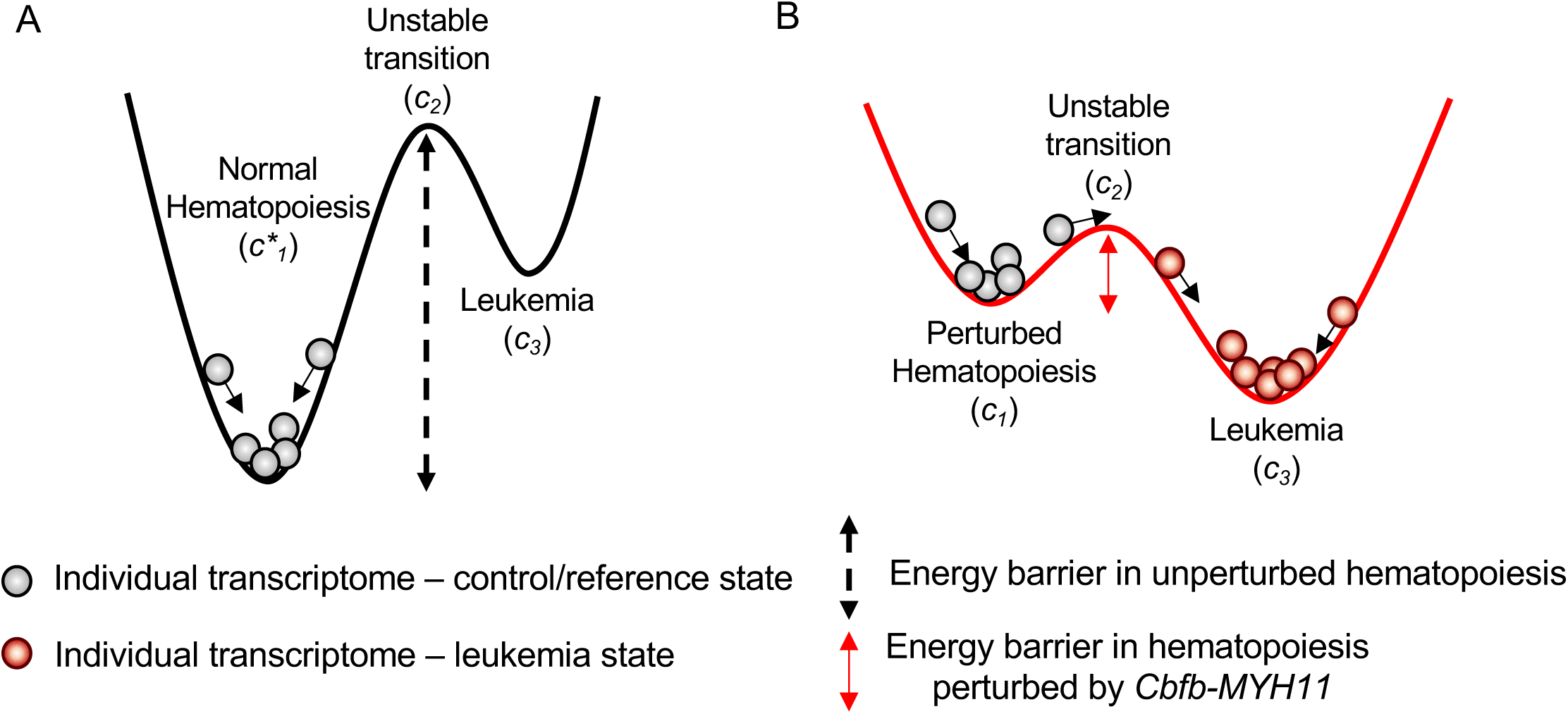
Cancer as a state-transition of the transcriptome. State-transition theory, the theory that is used to predict the transition of particles in a potential well, is applied to describe states of hematopoiesis, identify critical points in state-transition from health to leukemia, and to compute the probability of leukemia development. We model the action of oncogenic events as a reduction in the energy barrier required to cause state-transition, and thus increase the probability of cancer (leukemia) development. **A)** In unperturbed— normal—hematopoiesis, a large energy barrier between the reference state *c_1_* and unstable transition *c_2_*result in low probability of the state-transitioning to leukemia *c_3_*. **B)** In hematopoiesis perturbed by an AML oncogene *Cbfb-MYH11*, the energy barrier is reduced and therefore increases the probability of transition from *c_1_* to *c_3_* to a leukemia state. The * marker indicates normal hematopoiesis unperturbed by *Cbfb-MYH11*.

In this double-well quasi-potential, we postulated the existence of three critical points, denoted as *c*_1_, *c*_2_, *c*_3_. (Figure 1). The critical point 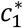 (* for control mice) represents a stable state of normal hematopoiesis whereas the critical point *c*_1_ represents a state of *CM*-perturbed hematopoiesis in *CM* mice (i.e., CM activation but not overt leukemia). The critical point *c*_2_ represents an unstable critical point that is a transition point in the dynamics from *c*_1_ to *c*_3_; *c*_3_ represents a state of overt leukemia corresponding to at least 20% circulating AML blasts. Because the critical point *c*_2_ is unstable, state-transition theory predicts that it would be unlikely to observe the system precisely at or very near this state. When the transcriptome-particle crosses the unstable critical point *c*_2_, the velocity of the particle increases. This can be interpreted biologically as an acceleration of the transcriptome-particle toward the leukemia state defined by the stable critical point *c*_3_. This prediction underscores the practical utility of state-transition theory in interpreting time-series genomic data. In fact, acquiring data precisely at each of the critical timepoints would be unlikely; therefore, theoretical constructs and mathematical predictions are necessary for contextualizing the relevance of data collected at any point or time near critical transitions.

To experimentally test our state-transition model based on transcriptome-particle dynamics from a state of normal hematopoiesis to a state of perturbed hematopoiesis and eventually to overt leukemia, we performed a longitudinal study using a well-established, conditional knock-in mouse model (*Cbfb^+/56M^/Mx1-Cre;* C57BL/6) that mimics a subset of human AML driven by the fusion gene *CM* (Figure 2A). In the conditional *CM* knock-in mice, CM expression is induced via the activation of Cre-mediated recombination by intravenous administration of synthetic double-stranded RNA polyinosinic–polycytidylic acid [poly (I:C)] (Supplemental Figure S1) (Cai et al., 2016; Kuo et al., 2006). Induction of CM expression results in perturbed hematopoiesis, lower energy barrier and increased probability of state-transition from normal hematopoiesis to leukemia. We collected PBMC samples from a cohort of *CM*-induced mice (n = 7) and similarly treated littermate control mice lacking the transgene (n = 7) before induction (t = 0) and at one-month intervals after induction up to 10 months (t = 1-10) or when the mouse was moribund. All but one of the *CM*-induced mice developed AML within the 10-month duration of the experiment (Figure 2B). The one remaining *CM* mouse exhibited CM-perturbed pre-leukemic expansion of progenitor populations in the bone marrow but leukemia had not yet manifested by the end of the 10-month study (data not shown). All PBMC samples were subjected to RNA-sequencing (Figure 2C) and flow cytometry analyses.

**Figure 2.**
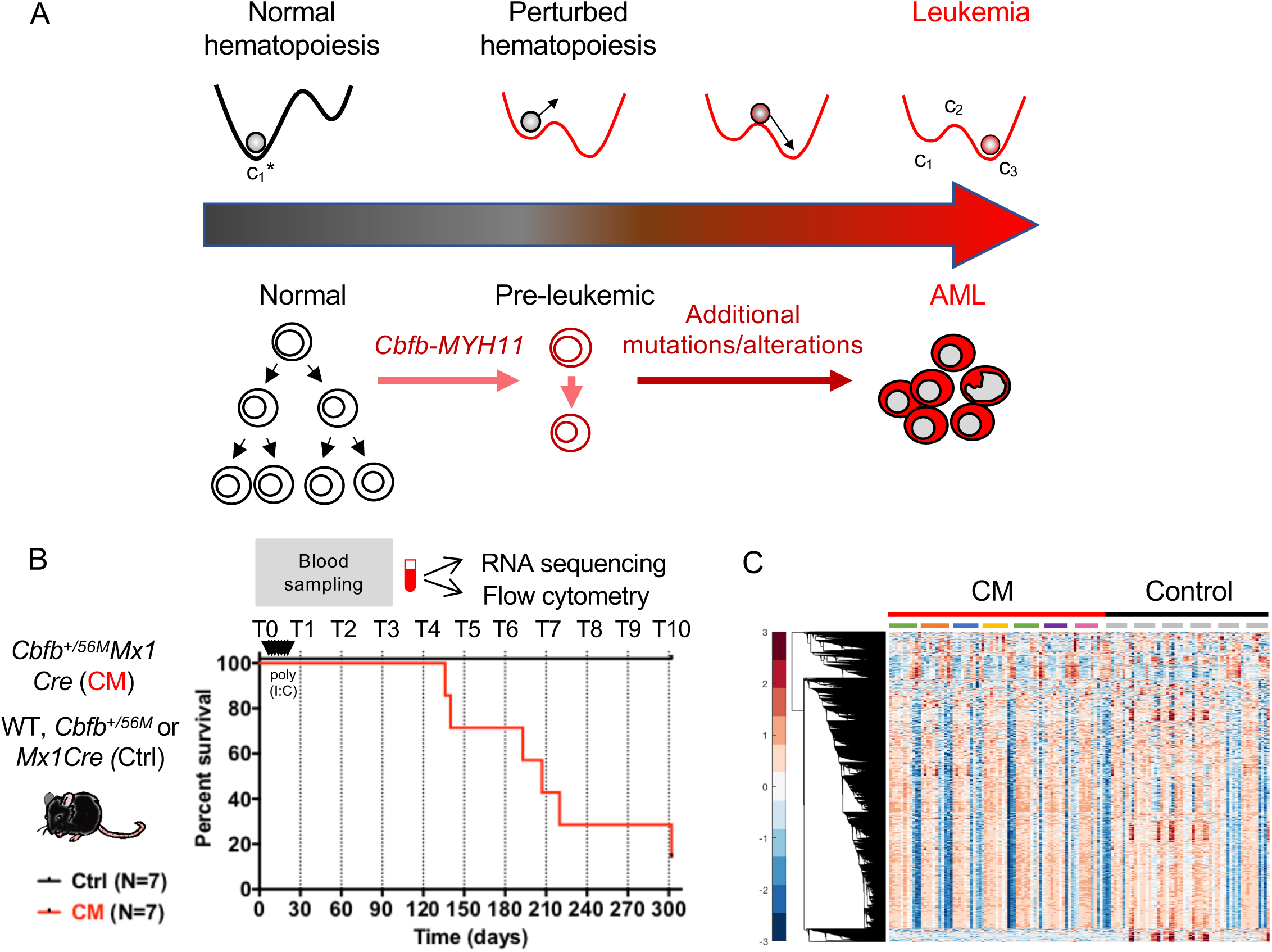
Experimental model of transcriptome state-transition. **A)** State-transition theory applied to leukemia development. The scheme represents the temporal evolution of the transcriptome from a healthy state to a leukemia state in a longitudinal study in the AML model induced by the *Cbfb-MYH11 (CM)* oncogene. In the conditional CM knock-in mouse model (*Cbfb^+/56M^Mx1Cre),* expression of *CM* in the adult bone marrow alters normal hematopoietic differentiation creating aberrant pre-leukemic progenitor cells which with time acquire additional genetic, epigenetic alterations needed for malignant transformation and AML development. **B)** In a cohort of CM mice (*Cbfb^+/56M^Mx1Cre*; N=7) and littermate controls (Ctrl; N=7) lacking one or both transgenes (*Cbfb^+/56M^ or Mx1Cre),* peripheral blood was sampled prior to and following CM induction (by poly (I:C) treatment) monthly for up to 10 months (or when mice were moribund with leukemia). Blood samples were subjected to bulk RNA-sequencing and flow cytometry analysis. Survival curve of CM (red line) and Ctrl (black line) mice corresponding to blood sampling time point (dashed line) are shown. **C**) Hierarchical clustering of time-series RNA-sequencing data for all samples (N= 132).

Given the positions of critical points, the double-well quasi-potential can be constructed up to a constant of integration by evaluating *U* = ∫ *U*′(*z*)*dz* where *U*′ = *dU/dx* = *α(x – c*_1_)(*x –c*_2_)(*x – c*_3_) where α is a scaling parameter and *x* is the spatial variable in the quasi-potential. The coefficients *α_i_* can be expressed in terms of *α*, *c*_1_, *c*_2_, *c*_3_ by expanding *U*′ and integrating with respect to *x*. State-transition theory predicts that the energy barrier—defined as the energetic difference between the initial state and the transition state—will be lowered by the expression of the *CM* gene, resulting in significantly increased probability and rate of transition from normal hematopoiesis to leukemia in *CM*-induced mice compared to control mice. In support of our hypothesis, we calculated α = 4.85 x 10^-8^ and observed that the energy barrier (*EB* ≡ *U(c*_2_) − *U(c*_1_)) is 0.99 (arbitrary units, A.U.) for *CM*-induced mice, nearly an order of magnitude lower than the energy barrier for control mice [6.45 (A.U.)].

### Construction of the leukemia transcriptome state-space

To represent the state-transition dynamics of the transcriptome from normal hematopoiesis to leukemia, we constructed a leukemia transcriptome state-space, in which the trajectory of the evolution from *CM*-perturbed hematopoiesis to development of overt AML could be geometrically represented. We initially performed genomic signal processing and dimension reduction analysis on the time-series RNA-sequencing data (Figure 2B). We constructed a matrix (*X*) such that each row corresponds to a sample and each column corresponds to a gene transcript level (log2 transformed counts per million reads)(Robinson et al., 2009). We then performed principal component analysis (PCA) to process and deconvolute the RNA-sequencing data into the components of the variance that most clearly associated with normal hematopoiesis or leukemia progression. Principal components (PCs) were computed via singular value decomposition (Figure 3A), which is a matrix factorization method. The singular value decomposition is computed on the mean-centered data 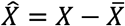 where 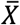 represents the column mean such that 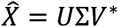 (* denotes the conjugate transpose, and *U* is a square matrix not to be confused with the quasi-potential energy polynomial). The columns of the unitary matrix *U* form an orthonormal basis for the sample space (i.e., the temporal dynamics of the transcriptome), the diagonal matrix Σ contains the singular values, and the columns of the matrix *V\** correspond to the eigengenes, or loadings, of each gene in the transcriptome per PC.

**Figure 3.**
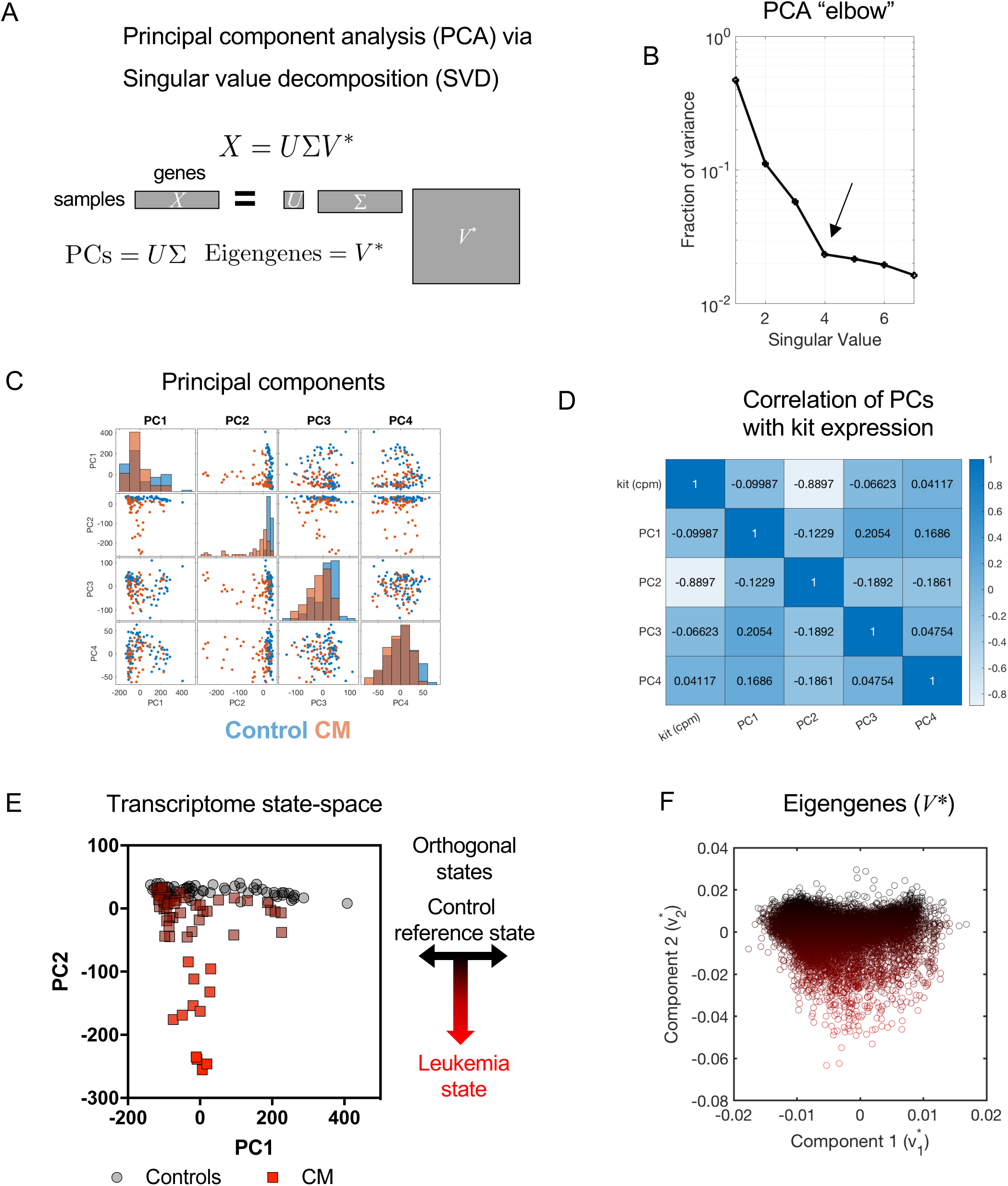
Construction of the transcriptome state-space. **A**) Singular value decomposition is used to factor the data matrix constructed with time-sequential (samples; rows) RNA-sequencing data (genes; columns) and perform principal component analysis (PCA). Principal components (PCs), singular values which reflect variance in the data, and gene weights (eigengenes) are computed. **B**) The fraction of the total variance in the data is plotted for the first seven singular values (all components shown in supplement Figure S2A). Black arrow indicates that PCA “elbow” at the 4^th^ singular value/component corresponding to 65% of the variance in the data. **C**) Each of the four first PCs plotted against each other (control: blue, CM: orange). Each dot is an individual transcriptome. Density of points along the x-axis is shown as a histogram on the diagonal. PC2 clearly shows the most separation between control and CM data (non-overlapping histograms). **D**) Correlation matrix of the first four PCs for CM samples with expression of *Kit,* a gene highly expressed and a surrogate phenotypic marker for leukemia cells. The second principal component (PC2) shows the strongest correlation with *Kit* (see also supplement Figure S3). **E**) PC2 encodes transition from control to leukemia. PC1 and PC2 create a 2D orthogonal transcriptome state-space. Points are individual transcriptomes from control and CM mice at different time points. **F**) The eigengene weights for the first two PCs are given by the first two columns of the matrix *V\** (shown in A) which may be associated with regions in the principal component state-space. The points in E and F are pseudo-colored from black to red, from north to south to indicate transition to leukemia.

We analyzed the singular values and 65% of the total variance was captured in the first 4 components, representing a majority of the variation in the data and corresponding to the PCA “elbow” (Figure 3B). A second elbow was identified at component 15 (see supplemental methods, Figure S2). A pairwise analysis of the first 4 components (Figure 3C) revealed that the second component (PC2) strongly correlated with the appearance of differentially expressed *Kit* (Figure 3D; supplemental Figure S4), which in this mouse model is a surrogate immunophenotypic marker for leukemic cells. We note that PCs are eigenvectors of the data matrix X, are orthogonal by construction and in fact the data projected along PC1 and PC2 have the smallest correlation coefficients (Figure 3D). We therefore chose PC1 and PC2 as orthogonal states (i.e. components) for the transcriptome state-space. We constructed a 2-dimensional state-space with the first (denoted as non-leukemic) and second (denoted as leukemic) sample components (*x*_1_, *x*_2_) = (*PC*1, *PC*2). We associated this space with the temporal dynamics of state-transition to leukemia so that the mean position of the reference (non-leukemic) state was located at PC2 = 0 and smoothly increased toward a leukemic state as the trajectory traveled south in the space (Figure 3E).

The second column of the loading matrix V* represents the eigengenes of leukemic progression (Figure 3F). Each gene is represented as a 2-dimensional vector which has components 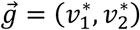. This representation enables the geometric decomposition of each gene into non-leukemic 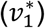 and leukemic 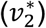 components, respectively (Figure 3F). This geometric analysis of eigengenes allowed us to interpret the leukemic component of genes based on their relative contribution to the leukemia state 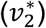 in differential expression analysis. We examined other dimension reduction methods to construct the state-space, but found them to be sub-optimal due to free parameters (e.g., diffusion mapping (Haghverdi et al., 2015)) or the inability to isolate leukemia trajectories with default settings (e.g., t-SNE (Pezzotti et al., 2016)) (see supplemental methods, Figure S5).

The construction of the state-space using PCA was not sensitive to variations in data-normalization method, sample number, or gene selection criteria, as shown by bootstrap cross-validation (see supplemental methods, Figure S6–S10). In fact, the geometry of the state-space could be inferred from time-series RNA-sequencing data derived from just 1 control mouse and 1 CM mouse (supplemental Figure S9) and was not changed by the exclusion of the known leukemia genes *Kit* or *CM* (supplemental Figure S10). These results demonstrate that PCA-based state-space construction is robust and reproducible regardless of variation in data-processing methods.

### Estimation of critical points in the transcriptome state-space

The PC-constructed transcriptome state-space allowed us to identify the position of the critical points in the state-space, to define the shape of the double-well quasi-potential and to predict the dynamics of the transcriptome-particle. We used K-means clustering to identify 3 clusters of the data along the leukemia axis (PC2) (Figure 4A). The centroid of all control samples was used as the point *c*_1,_*. The centroids of the CM clusters K1 and K3 were used as estimates of the leukemia critical points *c*_1,_ and *c*_3_, respectively. Because the critical point *c*_2_ is unstable, it is unlikely to be correctly identified with a centroid approach. To estimate *c*_2_, we constructed all quasi-potentials with values of *c*_2_ which ranged from to (Figure 4B, left), and computed the Boltzmann Ratio (BR) for each quasi-potential 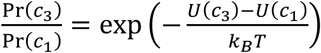 (Figure 4B, right). We assume the temperature is constant. We took *c*_2_ to be the “upper” boundary (maximum value) of the *c*_2_ cluster, which resulted in a BR of 81.4.

**Figure 4.**
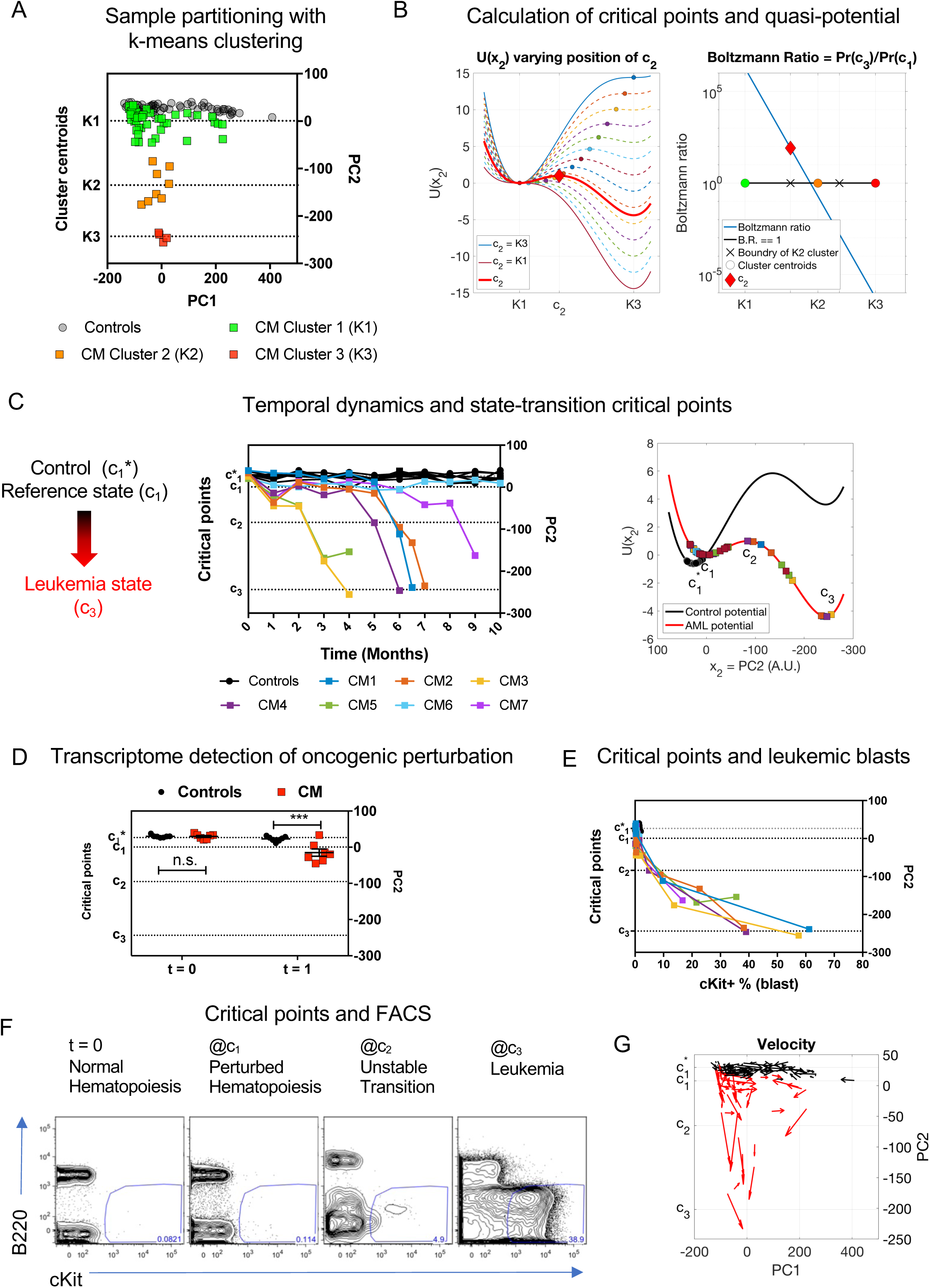
Critical points and state-space trajectory. **A**) The samples in the state-space for CM mice are associated with critical points *c_1_, c_2_*and *c*_3_ based on k-means clustering with k=3. The critical points *c_1_* and *c_3_*are the centroids of clusters 1 and 3 (*c_1_* = K1, *c_3_* = K3, respectively). **B**) Left: Because the transition critical point *c_2_* is unstable, we do not expect to observe the system precisely in this state. We therefore calculated the quasi-potential for a range of values of *c_2_* from *c_2_=c_1_* to *c_2_=c_3_*. Right: We calculated the Boltzmann ratio (B.R.=Pr(*c_3_*)/Pr(*c_1_*)) for quasi-potentials with the range of values of *c_2_*. Because the probability of transitioning to *c_3_* is greater than the probability of remaining at *c_1_* following CM induction (i.e. B.R.>1), we required that *c_2_* be chosen such that it is within the range of cluster 2 (indicated with “X” markers) and such that B.R. > 1. We therefore estimated the position of *c_2_* as the upper bound of cluster 2, resulting in a B.R. = 81.2 (red diamond, red curve). We also conducted a sensitivity analysis of this estimation on our subsequent model predictions (see supplementary methods). **C**) Temporal dynamics and state-transition critical points. Left: Transcriptome state-space trajectories of individual mice (controls in black; CM induced mice in colors). Right: state-transition critical points and dynamics of PC2 over time mapped onto the quasi-potential energy (*U(x)*) for control and CM mice. Controls remain at the reference state (*c*_1_*) and CM induced mice transition from the reference state of perturbed hematopoiesis (*c_1_*) to the leukemic state (*c_3_*). **D**) Early detection of oncogenic perturbation. The movement of transcriptome location in PC2 from *c*_1_* to *c_1_* can be detected at one month (t=1) after induction of CM expression (p < 0.01). **E**) The frequency of cKit^+^ leukemia blasts increases rapidly after crossing *c_2_* transition point. **F**) Representative flow cytometry plots of leukemia blasts (cKit^+^) frequency detected in the blood before induction and at each critical points. **G**) The velocity of movement in the state-space increases in magnitude after the critical point *c_2_* consistent with an acceleration of leukemia progression.

We then mapped the state-space trajectory of each mouse in time and the PC2 location relative to the estimated critical points *c*_1,_ *c*_2,_ *c*_3_ (Figure 4C; supplemental Figure S11). As early as one month (t = 1) following induction of CM (p<0.01) and despite the absence of any detectable circulating leukemic blasts, we were able to detect changes of the position in the transcriptome state-space (value of PC2) (Figure 4D), in line with the early hematopoietic perturbation driven by *CM* expression we previously reported(Cai et al., 2016; Kuo et al., 2009, 2006). Consistent with the prediction made using the state-transition theory, we observed acceleration of the transcriptome toward the leukemia state once the transcriptome-particle crossed the unstable critical point *c*_2_. Indeed, circulating leukemic blasts (cKit^+^) began to be detected (<5%) once the samples approached *c*_2_ (Figure 4E). After crossing this point, the acceleration toward the leukemia state was confirmed by a rapid increase of leukemic blasts and manifestation of overt disease (Figure 4F). The accelerated movement after crossing *c*_*_ was also supported by velocity calculated between each pair of time-sequential points in the 2D state-space (Figure 4G). Our results therefore support the applicability of the state-transition theory to cancer and demonstrate that leukemia development can be interpreted as transcriptome dynamics defined by critical points in the state-space trajectory.

### Prediction of leukemia progression

Because our model incorporates stochasticity that may be due to biological, experimental, technical, or time-sampling variations, the transcriptome trajectories cannot be precisely predicted by simply applying the equation of motion of the transcriptome-particle in the state-space double-well quasi-potential 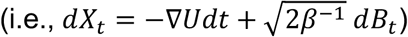. However, the stochastic model can provide examples of trajectories (Figure 5A). A well-established approach to characterizing and predicting stochastic dynamics is to consider the evolution of the probability density function. The spatial and temporal evolution of the probability density for the position of the particle *p(x*_2_, *t*) is given by the solution of the corresponding Fokker-Planck (FP) equation

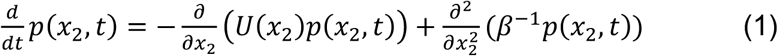

where *β*^−1^ is the diffusion coefficient, which may be estimated with a mean-squared displacement analysis of particle trajectories (see supplemental methods, Figure S11)(Pavliotis, 2014). Solution of the FP permits the direct calculation of the expected first arrival time from an initial point (i.e., perturbed hematopoiesis) in the state-space to a final point (i.e., leukemia). In our study, we calculated the mean time to develop leukemia for the cohort of mice by numerically solving the FP equation with the initial condition taken from the data (Eq. 1, Figure 5B-C). We calculate the probability of being in the leukemia state at time *t = t_i_* from a given initial position in time and space by integrating the probability density past *c_2_* from time *t = 0* as 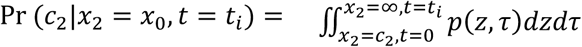. The mathematical model accurately predicted both the proportion of induced mice that will develop leukemia as well as the mean time for disease to manifest; in fact, the predicted and actual median survival time, defined as the median time to develop leukemia (at least 20% circulating blasts), were not significantly different (p >0.05) (Figure 5D).

**Figure 5.**
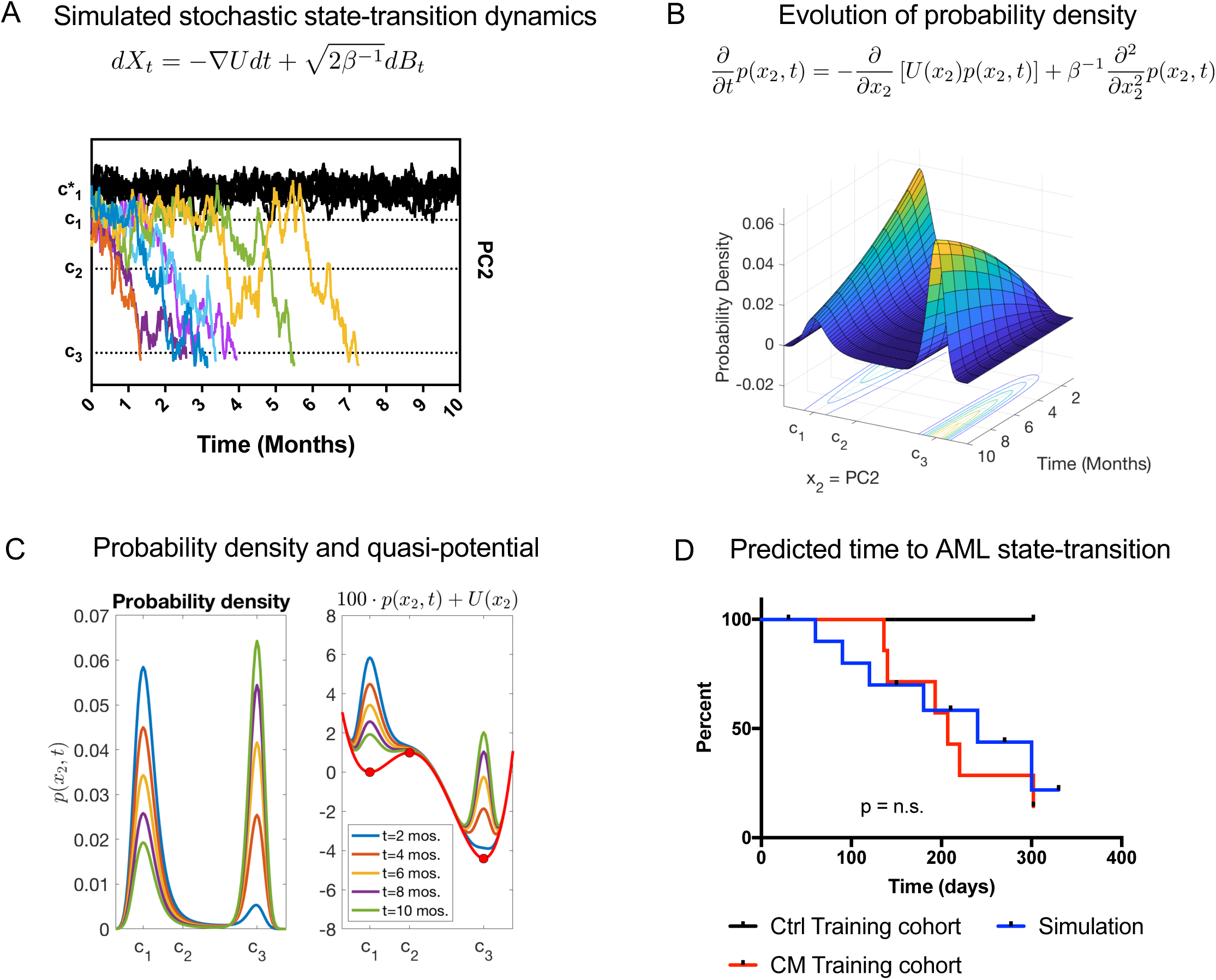
Theory and predictions of state-transition dynamics. **A**) The equation of motion in the quasi-potential is a stochastic differential equation which predicts trajectories of state-transition. One realization of a stochastic simulation is shown as trajectories of the transcriptome particle in state-space (PC2) over time (controls black, CM induced mice in colors; colors are matched with observed trajectories for CM mice as shown in Figure 4B-C). Controls remain at the reference state (c*_1_) and CM induced mice transition from the reference state of perturbed hematopoiesis (c_1_) to the leukemic state (c_3_). **B**) Due to the stochastic nature of the biological process and variability in RNA-seq data, we predict state-transition by considering the spatial-temporal evolution of the probability density (*p(x,t)*) given by numerically solving the Fokker-Planck equation. **C**) Selected time solutions of the FP shown in B (left) overlaid with the CM quasi-potential *U*(*x*_2_) to illustrate the transition of probability density from the first state to the second state corresponding to the wells of the quasi-potential, respectively (right). **D**) The predicted (simulated) time to state-transition is calculated with with the probability density equation and compared to observed time to leukemia with a survival analysis. The observed and simulated time to AML are not statistically different from each other (p>0.05) n.s., not significant.

### Biological interpretation of the critical points in the transcriptome state-space

Because state-transition theory enables the interpretation of time-series genomic data in terms of critical points, we hypothesized that the transcriptional events [differentially expressed genes (DEGs)] occurring at each critical point (*c*_+_, *c*_*_, *c*_)_) represented critical biological alterations driving the transition from normal hematopoiesis to perturbed hematopoiesis and eventually to leukemia (Figure 6A). To identify these events, we partitioned the data such that each sample was associated with a unique critical point by calculating the nearest proximity of a given sample to the location of the critical points in the state-space (Figure 4A). We then identified DEGs by performing pairwise comparisons of gene expression at each of the critical points with that of the reference state (i.e., *c*_1_ vs 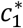; *c*_2_ vs 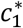; *c*_3_ vs 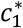) and with other critical points (*c*_2_ vs *c*_1_; *c*_3_ vs *c*_1_; *c*_3_ vs *c*_2_) using edgeR and a false discovery rate of 0.05 (supplemental Figure S13). The number of DEGs (defined as q<0.05, |log2(FC)|>1) for each comparison is summarized in Table 2 (see supplemental Table S1-6 for gene list). We then categorized the DEGs at each critical point as early events (*c*_1_ vs 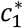, not in *c*_2_ vs *c*_1_ or *c*_3_ vs *c*_2_), transitional events (*c*_2_ vs 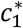, not in *c*_2_ vs 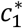 or *c*_3_ vs *c*_2_), and persistent events (genes that remained as DEGs at all three of the critical points; *c*_1_ vs 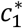; *c*_2_ vs 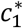; *c*_3_ vs 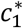) (Figure 6A).

**Figure 6.**
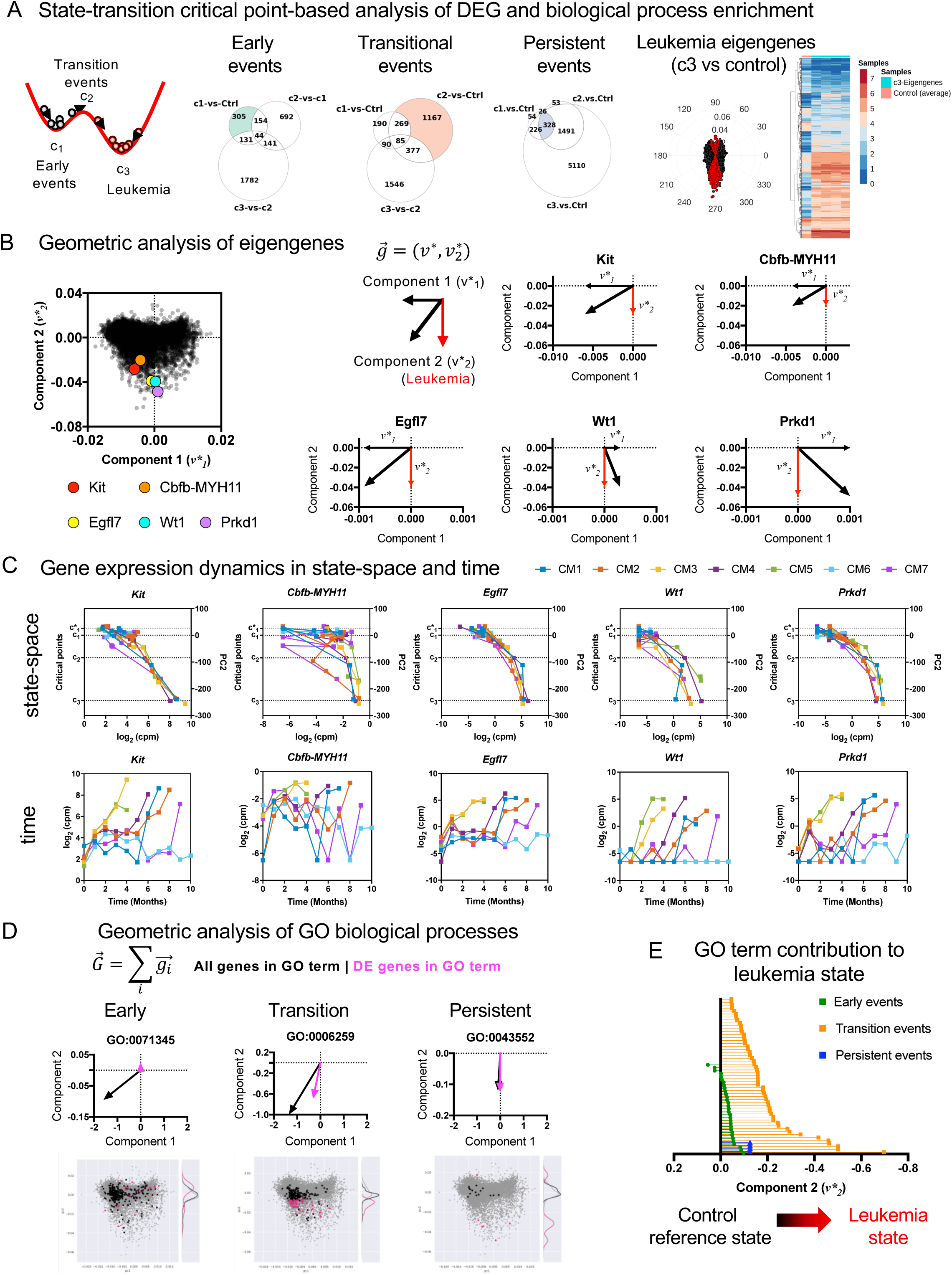
State-transition based analysis of genes and pathways in leukemia progression. **A**) The state-transition theory is used to group samples relative to critical points. Early, transition, and persistent events in leukemia progression are defined relative to critical points. Gene expression of each differentially expressed gene in a single GO term with the smallest q-value in the subset analysis is shown for each subset analysis (early, transition, and persistent events) relative to critical points. **B**) Geometric representation and decomposition of *Kit, CM, Egfl7*, *Wt1*, and *Prkd1,* demonstrating their contribution to leukemia state. Eigengene weights (*V\**, see Figure 2A,F) for the first two components are plotted for all genes (black). Selected genes (*Kit, CM, Egfl7*, *Wt1*, *Prkd1*) are shown in colors and are oriented south in the space indicating a contribution to leukemia. **C**) Genes expression dynamics in state-space and time. Expression levels for selected leukemia eigengenes *Kit, CM, Egfl7*, *Wt1*, *Prkd1* are plotted against location in the leukemia (PC2) state-space (top) or time (bottom) for each CM leukemia mouse. Increasing expression of these genes are concordantly correlated with movement from normal hematopoiesis (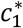) to leukemia (*c*_1_) in the state-space. **D**) Top: Geometric and vector analysis of GO terms overall (black) and genes (pink) that are differentially expressed (DE). The sum total of genes in a GO term (black vector) and the portion which is differentially expressed (pink vector) are used to geometrically interpret the contribution of the GO term towards leukemia progression. Bottom: The eigengene state-space is used to represent genes and biological pathways identified through pathway enrichment analysis as vectors. Subsets of genes in a given pathway which are differentially expressed are shown in pink and the distribution along the second component is shown as a kernel density. **E**) The leukemia component of the vector representation of each GO term enriched in the early, transition and persistent events shown. Biological pathways represented as vectors demonstrate increasing orientation in the state-space towards the state of leukemia (*v*_2_*).

**Table 1.** Glossary of terms

| Term | Meaning |
| --- | --- |
| State variable | A state variable is one of a minimal set of variables that describe the mathematical "state" of a dynamic system. In this work, the state variable is the transcriptome derived from RNA-sequencing of peripheral blood mononuclear cells. |
| State-space | A state-space is a mathematical representation of all possible configurations of a system defined by the state variables. In this work, the state-space is constructed with principal component analysis of time-series RNA-sequencing data of peripheral blood mononuclear cells over the course of leukemia progression in a mouse model. |
| State-transition | A state-transition is the dynamic process of a system changing from one state to another. In this work, the state-transition of interest is the transition of the transcriptome from a reference state of hematopoiesis to leukemia. |
| Probability density | The probability density gives the probability of finding the system in a given state (position in the state-space) at a given time. The probability density takes values between zero and one. The sum of the probability density over the entire state-space is one. The probability density is given by the solution of the Fokker-Planck equation. |
| Double-well quasi-potential | A double-well potential is an energy function which has two local minima, similar to a "w" shape. In this work, the double-well potential is derived from the transcriptome state-space. The potential is referred to as a "quasi-potential" because the state-space does not have physical units, and therefore the potential energy function does not have a clearly defined physical analog. The wells of the potential correspond to states of the transcriptome. |
| Eigengene | Eigengene refers to the coefficient weights (or loadings) of a given gene computed with principal component analysis. In this work, a "leukemia eigengene" refers to the weight of a given gene in the principal component analysis of gene expression data associated with leukemia. |

**Table 2.** Differentially expressed genes based on critical points. Thousands of genes are differentially expressed across critical points in the transcriptome state-transition (q<0.05, |log2(FC)|>1).

| Test vs. reference | #genes down | #genes up | #genes total |
| --- | --- | --- | --- |
| **c<sub>1</sub> vs control (c<sub>1</sub><sup>\*</sup>)** | 2,305 | 1,859 | 4,164 |
| **c<sub>2</sub> vs control (c<sub>1</sub><sup>\*</sup>)** | 4,421 | 4,126 | 8,547 |
| **c<sub>3</sub> vs control (c<sub>1</sub><sup>\*</sup>)** | 6,119 | 5,515 | 11,634 |
| **c<sub>2</sub> vs c<sub>1</sub>** | 3,560 | 3,772 | 7,332 |
| **c<sub>3</sub> vs c<sub>1</sub>** | 5,744 | 5,565 | 11,309 |
| **c<sub>3</sub> vs c<sub>2</sub>** | 3,602 | 3,274 | 6,876 |

Gene Ontology (GO) analysis revealed insights into the biological and functional impact of DEGs identified on the basis of critical points in the transition from normal hematopoiesis to leukemia (supplemental Table S7-9). For early transcriptional events at *c_1_*, the top three GO terms ranked by q-value (multiple-test corrected p-value) included extracellular matrix organization (GO-0030198), cellular response to cytokine stimulus (GO-0071345) and cytokine-mediated signaling pathway (GO-0070098) (Figure 6A; Figure S14A; Table S7). For the transitional events at *c_2_*, the top three ranked GO terms included DNA metabolic processes (GO-0006259), DNA replication (GO-0006260), and G1-S transition of mitotic cell cycle (GO-0000082) (Figure 6A; Figure S14A; Table S8). DEGs occurring at *c_1_*, which continued to be differentially expressed at the critical points *c_2_*and *c_3_*, were defined as persistent events. Consistent with cKit expression marking the leukemia blasts, *Kit* up-regulation was among the persistent events seen at *c_1_, c_2_,* and *c_3_*. For persistent events, the top three ranked GO terms included positive regulation of phosphatidylinositol 3-kinase activity (GO-0043552), positive regulation of phospholipid metabolic process (GO-1903727), and positive regulation of lipid kinase activity (GO-0090218) (Figure 6A; Figure S14A; Table S9) We note that hierarchical clustering correctly identified leukemia samples but failed to differentiate CM from controls at intermediate time points or to partition the samples temporally (supplemental Figure S12).

Given the 2-dimensional geometry of the transcriptome state-space, we were also able to decompose the contribution of each gene to leukemia progression by considering the second component of the eigengene vector 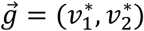. For instance, considering the expression of *Kit* and *CM* genes, we showed that the magnitude of the second component of both genes was negative (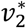 < 0) with 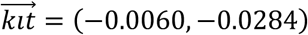 and 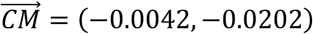, confirming their contribution to promote leukemia development (Figure 6B). Notably, the proangiogenic factor *Egfl7* 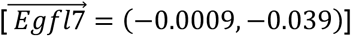 (Papaioannou et al., 2017), leukemia-associated antigen *Wt1* 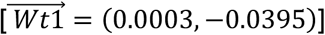 (Adnan-Awad et al., 2019; Chaichana et al., 2009; Løvvik Juul-Dam et al., 2019) and the protein kinase *Prkd1* 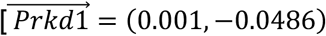 genes are among those showing a strong selectivity in the direction toward leukemia in the eigengene space (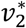 < 0) (Figure 6B). Consistent with a positive contribution promoting leukemia progression, all CM mice that developed leukemia (CM1-5, CM7) showed increasing expression of these leukemia eigengenes (*Kit*, *CM*, *Egfl7*, *Wt1*, *Prkd1*) and reproducible trajectories as they move from normal hematopoiesis (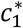) to leukemia (*c*_3_) (Figure 6C). The trajectories of these leukemia eigengenes in the state-space were remarkablly concordant for all CM leukemia mice (Figure 6C, top), in constrast to the nonsynchronous trajectories observed in time (Figure 6C, bottom). Therefore, geometric analysis through the transcriptome state-space provides a robust and meaningful approach to identity relevant gene expression signals and to examine leukemogenic contribution of individual gene as well as the collective contribution of a set of genes.

With this geometric interpretation in hand, we quantified each GO term as the vector sum of each constituent eigengene, so that 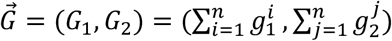. The contribution of an individual GO term to the progression from normal hematopoiesis state to a leukemia state was also given by the second component of the summed vector, *G*_2_, which represented the maximum contribution of an individual GO term pathway *G* to leukemogenesis (Figure 6D; black vector). Because not all of the constituent genes for a GO term (black dots) were differentially expressed (pink dots), we considered only the subset of genes in the GO term that actively contributed to leukemogenesis as evidenced by differential expression (Figure 6D; pink vector). As such, each significantly enriched GO term could be quantitatively analyzed for its relative contribution to the development of leukemia as the sum total 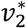 contributions of the DEGs in that particular GO term.

To evaluate the potential leukemogenic contribution of of the GO terms corresponding to the step-wise perturbation, we also performed vector analysis of each GO pathway enriched in early, transitional, and persistent events and represented them as vectors in the state-space (Figure 6D). Notably, our analysis of *c*_+_ early events revealed that some highly significantly enriched pathways which exhibited contributions away from leukemia (north, 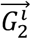 > 0)(Figure 6E; Figure S15), suggesting the presence of a restorative force in initially CM perturbed hematopoiesis, which attempts to revert the perturbed system back to the reference state of normal hematopoiesis. On the other hand, analysis of GO terms enriched for transitional and persistent events demonstrated an increasing magnitude and direction (angle) toward the leukemic state (Figure 6D, Figure S16-18). Evaluation of 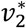 for all early, transitional, and persistent GO terms revealed a strong overall leukemogenic contribution given by transition events occurring at *c*_2_ (Figure 6E), underscoring the unique biological insights gained by differential analysis based on state-space critical points.

Analysis of the leukemic transcriptome at *c*_3_ showed dysregulation of a large number (11,634) of genes (supplemental Figure S15B), making it difficult to perform pathway enrichment or to interpret in terms of contribution to leukemia. Thus, we filtered genes based on the angle subtended (*θ*) by the gene in the 2-dimensional state-space. The range of angles (*θ*) identified as being associated with leukemia evolution was defined as the minimal sector of a circle centered at (0,0) that included all leukemic samples as well as the mirror image of this sector along the absicca axis of symmetry (supplemental Figure S19). This selection identified differentially expressed leukemia eigengenes (leukemia eigenDEG). The top three GO terms for leukemia eigenDEG ranked by q-value (multiple-test corrected p-value) included mitotic spindle organization (GO-0007052), centromere complex assembly (GO-0034508), and microtubule cytoskeleton organization involved in mitosis (GO-1902850) (Table S10), consistent with the hyper-proliferative phenotype, leukemic cell trafficking, and extramedullary tissue infiltration associated with late-stage disease.

Altogether, our state-transition model, leukemia transcriptome state-space, and leukemia eigengenes offer biologically relevant interpretation and insights into the complex multistep process of malignant transformation and disease evolution.

### Validation study with independent cohorts

To validate our state-transition mathematical model, state-space, and analytical approach, we collected PBMC RNA-sequencing data from two additional independent cohorts of control and CM mice. We collected validation (v) cohort 1 samples monthly for up to 6 months; (vControl1-7; vCM1-9) and collected validation cohort 2 samples sparsely at 3 randomly selected timepoints; (vControl8-9; vCM10-12) during leukemia progression. We performed PCA analysis of the validation cohort 1 and 2 data, which again demonstrated that the majority of the variance was encoded in the first 4 PCs (supplemental Figure S20A) and the leukemia-related variance was encoded in PC2 (Figure 7A). We then evaluated our ability to map state-transition trajectories and predict leukemia development in the validation cohorts by projecting the data from the validation cohorts into the state-space constructed using the training cohort. We accomplished this by using the eigengenes from the singular value decomposition of the training data as follows: a matrix of new data, 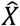, was projected into the state-space by multiplying 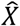 by the matrix *V* from the training data, so that the coordinates of the new data in the reference state-space were given by *PC* =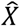*V*. Because the matrices 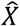 and *V* must have the same dimension, and more importantly the weights of the genes in the matrix *V* must match one-to-one with the genes in 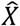, the projection may use only genes in the intersection of *X* and 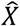.

**Figure 7.**
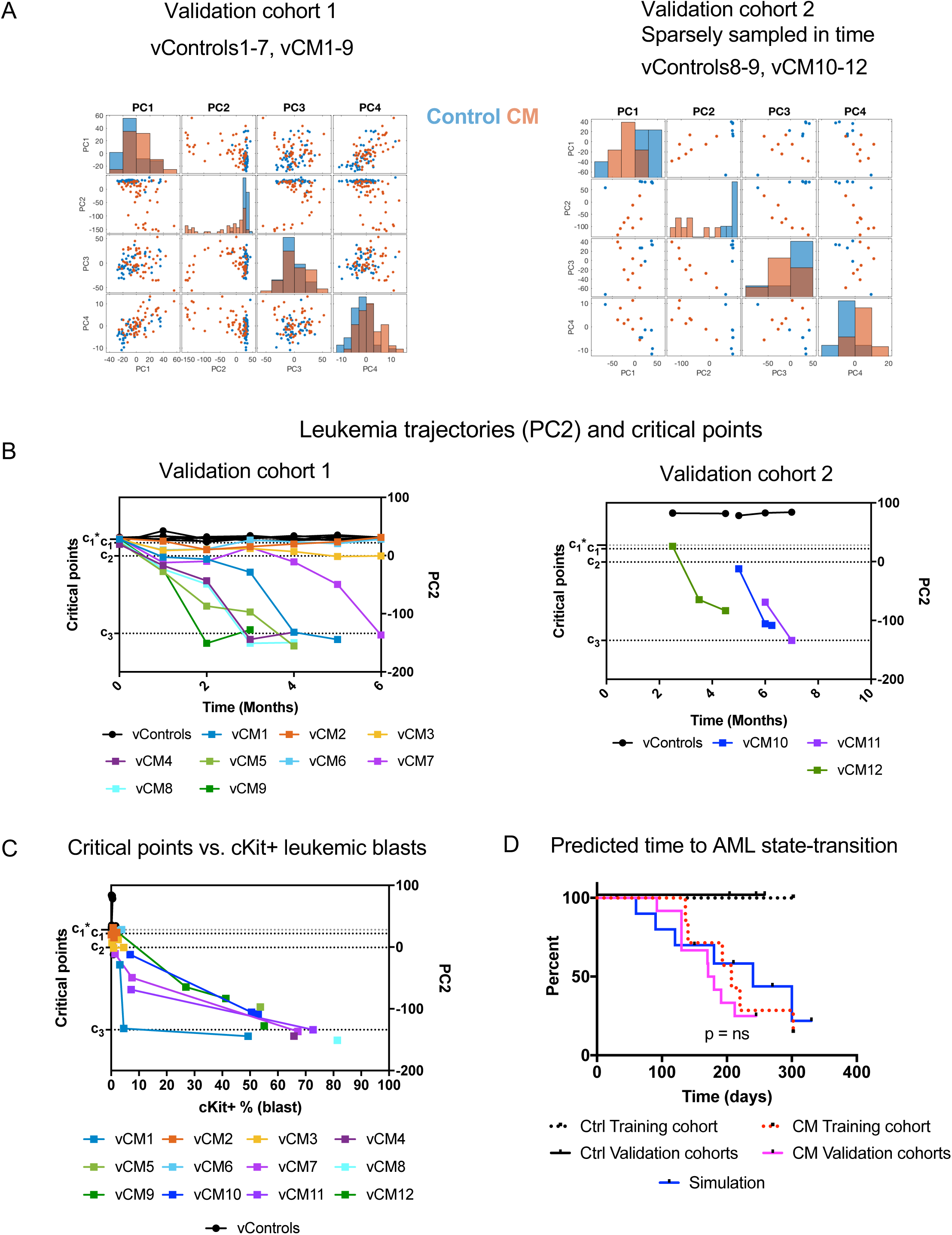
Validation cohorts. As a validation of the eigengenes, mathematical model, critical points, and state-space, we mapped data from two additional independent experiments: cohort 1 and cohort 2. Cohort 1 consists of 7 controls and 9 CM mice sampled at the same frequency as the training cohort. Cohort 2 consists of 2 controls and 3 CM mice sampled sparsely in time. **A**) The first four components of PCA for the validation cohort data are plotted against each other. The density of data points along the x-axis of each PC is shown with histograms on the diagonal. The pairwise plots clearly indicate that CM samples and leukemia trajectories are encoded in PC2. This can be seen by the separation of the control and CM distributions in grid cell (2,2), as compared to overlapping distributions in the other components. **B**) Leukemia trajectories (PC2) of validation cohorts projected into the state-space constructed with the training cohort. Critical points were estimated with the same procedure as the training data set. **C**) The frequency of cKit^+^ leukemia blasts increases rapidly after crossing c_2_ transition point. **D**) The predicted (simulated) time to state-transition is calculated with the probability density equation and compared to observed time to leukemia with a survival analysis for validation cohorts (cohort 1 and 2; CM n=12; Ctrl n=9) The observed and simulated time to AML are not statistically different from each other (p > 0.05). n.s., not significant.

Thus, we mapped the trajectory of each mouse in the validation cohorts in the PC2 space (Figure 7B). State-transition critical points *c*_1,_ *c*_2,_ *c*_3_ were estimated using k-means clustering (k=3), using the same method that was used on the training cohort (Figure 4A-B). The trajectories of vControls remained at *c*_1,_*, and vCM mice progressed toward a leukemic state at *c*_3_, as in the original analysis. We detected CM mice at *c*_1_, 1 month (t = 1) after induction of CM expression (supplemental data Figure S20B-C), similar to the initial dataset (Figure 4D). We observed similar state-space trajectories, in that we detected acceleration of the transcriptome toward the leukemia state once it crossed the unstable critical point *c*_2_ (Figure 7C).

We solved the FP equation for probability density (Eq. 1) with initial conditions derived from the validation cohort data and parameters estimated from the training cohort, which accurately predicted the time to leukemia for all CM mice (n=9) that developed leukemia during the study period (p>0.05, Figure 7D). Three CM-induced mice in validation cohort 1 (vCM2, 3, 6) did not develop leukemia during the 6-month study period, and were mapped to positions in the state-space between *c*_1_ and *c*_2_ but did not cross the transition point *c*_2_ (Figure 7B), consistent with a delayed leukemia onset predicted by our model. These mice showed pre-leukemic states in the bone marrow (i.e., expansion of pre-leukemic progenitor populations) at the end of the study, indicating leukemia progression was taking place but had not yet manifested (see supplemental data Figure S21). Altogether, these results demonstrate our ability to reproduce state-transition trajectories, identify critical points and predict leukemia development in multiple independent cohorts. Thus, our state-transition mathematical model, transcriptome state-space, and analytical approach provides a robust framework to contextualize and interpret global and temporal changes in gene expression, and use them to predict system dynamics relevant to leukemia development.

## Discussion

Here we report the application of the theory and mathematics of state-transitions to interpret temporal genomic data and accurately predict AML development. As a proof of principle, we used time-sequential RNA-sequencing data from a well-characterized orthotopic mouse model of inv(16) AML and applied a mathematical model of state-transition from health to leukemia. We demonstrate the feasibility of predicting the trajectory of state dynamics and time to leukemia using time-sequential genomic data collected at sparse timepoints.

Recent studies have shown that somatic mutations may precede diagnosis of acute myeloid leukemia (AML) by months or years(Desai et al., 2018) and that deep sequencing of mutations can be used to differentiate age-related clonal hematopoiesis from pre-AML and predict AML risk in otherwise healthy individuals(Abelson et al., 2018). However, these approaches rely on population statistical analysis and do not provide a hypothesis-based theory which can predict specific time to leukemia development in individual patients or be translated to other cancer settings. One of the greatest challenges in analyzing time-sequential data is the fact that the signal(s) of interest (i.e., leukemia) often are not synchronized in time. Although methods exist to perform time realignment, designed to overcome this issue (Liu and Müller, 2003; Liu and Yang, 2009; Sun et al., 2016, 2008) they require prior knowledge of the system dynamics. Here, we implemented a method that does not require a priori information, and allows for any nonlinear time dynamics, including transient changes to elements of the eigengene due to stochastic variations or biological fluctuations, including environmental conditions that may be random. Furthermore, our approach guides interpretation of temporal genomic data even when data are incomplete or sparse—as is often the case with longitudinal data from the clinic.

We constructed the leukemia transcriptome state-space based on PCA of RNA-sequencing data obtained from the peripheral blood over the course of leukemia progression. PCA is most often employed to study variance in a dataset and to cluster samples with similar levels of variance. When the PCs can be interpreted, the method provides both a means of reducing the dimensionality of the dataset and also a means to gain insight into the data. In our case, PCA yielded insight into the temporal dynamics of leukemia progression that are encoded in a single component (PC2). The corresponding transcriptome state-space allowed us to study leukemia dynamics by isolating the transcriptional events directly affecting the transition to a leukemia state. Using a mathematical model of state-transition, we demonstrated our ability to predict the time to develop leukemia from a state of perturbed hematopoiesis based on changes in the transcriptome of circulating blood cells over time.

Our results show that movement through the transcriptome state-space can be understood in terms of critical points—mathematically-derived inflection points—which provide a framework to predict the development of leukemia for any point in the space, at any timepoint in leukemia progression. The double-well quasi-potential mathematical model allows us to compute the probability of transition from one state to another for any point in the state-space by numerically solving the FP equation (Figure 5C). This approach may be generalized for any two-state system for which a distance measured between the states can be defined. The trajectories and dynamics in the state-space and the theory of state-transition provided disease-relevant context to guide our analysis of complex time-sequential gene expression data.

Through the analysis of DEGs at each critical point and by categorizing early, transitional, and persistent events according to GO classification, we identified critical step-wise transcriptomic perturbations during leukemogenesis. Early events are enriched for cellular response to cytokine stimulus and cytokine-mediated signaling pathway, consistent with previously reported altered cell signaling and impaired lineage differentiation induced by the *CM* oncogene (Cai et al., 2016; Kuo et al., 2009). Notably, our results revealed that early perturbations associated with critical point *c*1 are not necessarily contributing positively to leukemogenesis but may instead represent counteracting homeostatic response. In addition, the unstable critical point *c*2 was characterized by aberrant expression of many genes involved in DNA damage and DNA repair, consistent with the notion that additional cooperating mutations or epigenetic alterations are necessary for full leukemia development (Cai et al., 2016; Kuo et al., 2006). Furthermore, we identified genes that, although not uniquely associated to individual critical points, were persistently and differentially expressed at all critical points *c_1_, c_2_,* and *c_3_* during the leukemia state-transition. These genes are mainly involved in signaling pathways that support cell proliferation and survival, and vector analyses demonstrated a direction of strong contribution toward PC2 (leukemia). These persistent events can be interpreted as a force cooperating with the *CM* oncogene to propel the movement of the transcriptome-particle from the reference state to the leukemia state.

Thus, in addition to the mathematical and physical interpretation of state-transition dynamics, our approach provides a possible biological connotation for the critical points identified by the state-transition theory. Furthermore, the location and trajectory of individual genes in the state-space allows assessment of the direction and the magnitude of leukemia contribution. For example, *Egfl7*, *Wt1* and *Prkd1* showed a strong selectivity in the direction toward leukemia and their expression level consistently increased during transition toward leukemia, particularly between *c_1_* and *c_2_* in the state-space. Unlike the nonsynchronouse changes for individual CM leukemia mouse observed in time, these leukemia eigengenes displayed consistenly concordant trajectories in the leukemia state-space. These results reaffirm that our transcriptome state-space and approach indeed can infer information relevant to the phenotype of interest (i.e., leukemia), and highlight the utility of location in state-space to guide analysis and biological interpretation of temporal data. Notably, *Egfl7*, *Wt1* and *Prkd1* are among the persistent differentially expressed genes at all critical points *c_1_, c_2_,* and *c_3_* during leukemia state-transition. *Egfl7* is known to be highly expressed and predict poor prognosis in AML(Papaioannou et al., 2017), and is also a host gene of *miR-126*, which is a miRNA signature associated with inv(16) AML(Li et al., 2008). Overexpression of *WT1* has been found to predict poor prognosis and minimal residual disease in AML(Adnan-Awad et al., 2019; Becker et al., 2010; Løvvik Juul-Dam et al., 2019). *Prkd1* encodes a serine/threonine protein kinase and is part of all top 3 ranked GO terms enriched for persistent events. The specific role of *PRKD1* in AML has not been described, however, it is known to promote invasion, cancer stemness and drug resistence in several solid tumors(Döppler et al., 2016; Kim et al., 2016; Zhang et al., 2018). These results highlight the relevance of biological insights provided by the geometry of the transcriptome state-space and critical points of state-transition theory.

State-transition theory and corresponding mathematical models have been applied to other systems and to other omics data platforms (e.g., epigenomics, miRomics)(Zadran et al., 2014, 2013; Zhou et al., 2012). However, our application of state-transition theory to the interpretation of cancer evolution is novel not only because is applied to time-series data, but also because of the use of bulk RNA-sequencing data collected from the peripheral blood. Notably, this approach allows us to derive useful information about the state of the system as a whole, without concern for the heterogeneity related to additional mutations, clonal dynamics, or composition of cells within the sample. Given that additional mutations, epigenetic alterations, and clonal heterogeneity converge into gene expression profiles collectively representing perturbations that are ultimately responsible for the observed phenotypes, our use of RNA expression profiles represented as configurations of the system allows integration of informative quantities relevant to the disease state, without suffering from a combinatorial explosion of complexity. We showed that hierarchical clustering was unable to distinguish CM samples from controls or to partition the samples temporally. Other methods currently available to analyze time-series genomic data (Fischer et al., 2018; Spies et al., 2017; Spies and Ciaudo, 2015; Sun et al., 2016) were not able to: 1) predict the time to develop leukemia, 2) predict the effects of changes in sets of genes over time, or 3) quantify the relative importance of one gene set or pathway over another. Other approaches (i.e., surprisal analysis) that use concepts of thermodynamic (non)equilibrium and statistical mechanics(Facciotti, 2013) are also useful tools for analyzing cellular transitions (e.g., epithelial to mesenchymal transition(Zadran et al., 2014) and early stages of carcinogenesis(Remacle et al., 2010)), but to our knowledge they have not been used to analyze temporal data and do not provide similar geometric- or critical point-based interpretation of the data.

Although data from a relatively simple mouse model of AML were used to develop this theoretical framework, we demonstrated that the results are reproducible in multiple animal cohorts (i.e., one training cohort and two validation cohorts) and that the robustness of this approach is not affected by variability in sampling time, frequency, sample preparation, or data normalization methods (supplemental Figure S6–S10). Future studies will include prospectively collected time-sequential samples from healthy donors and patients with leukemia. Furthermore, it would be informative to determine the extent to which AML driven by different oncogenes share biological connotations and interpretations derived from the perspective of state-transition critical points. These information will guild further refinement and application of the framework to analyze diverse patient samples in the clinic.

We show that early perturbation could be detected based on the transcriptome state-space and early transcriptome trajectory could predict disease development using our mathematical model of state-transition. In the future, our state-transition dynamical model could be applied in the clinic to support proactive monitoring that detects early perturbations away from a reference state in a patient/individual over time, adding a mathematical and theoretical component to recent reports of longitudinal deep-sequencing to monitor changes in health over time(Schüssler-Fiorenza Rose et al., 2019). The reference state could be a healthy state prior to disease or could be a diseased state, in which case the orthogonal direction would be interpreted as transition back to a state of health upon therapeutic intervention. To this end, possible applications include monitoring of patients who achieved complete remission post-treatment and are closely followed for relapse, or patients with clonal hematopoiesis of indeterminate potential (CHIP) undergoing chemo or radiation therapy for unrelated cancer and who are at high risk of developing therapy related myeloid neoplasms(Bowman et al., 2018; Genovese et al., 2014; Sperling et al., 2017). This approach could be also applied to evaluation of antileukemia therapies, and to predict achievement and duration of treatment response at early time points. Therefore, our state-transition analysis approach has potential applications for prospective monitoring in disease prevention, early detection, and predicting response to therapy and disease relapse not only in leukemia but also in other types of cancer.

## Acknowledgments

This work was supported in part by the National Institutes of Health under award number R01CA178387 (to Y.-H.K) and R01CA205247 (to Y.-H.K/G.M.) and the Gehr Family Center for Leukemia Research. Research reported in this publication included work performed in the Integrated Genomics Core, Bioinformatics Core, Analytical Cytometry Core, and Animal Resource Center supported by the National Cancer Institute of the National Institutes of Health under award number P30CA33572. The content is solely the responsibility of the authors and does not necessarily represent the official views of the National Institutes of Health.

## Author Contributions

Conceptulization, R.C.R, S.B., Y-H.K and G.M.; Methodology, R.C.R, S.B., Y-H.K, D.F, D.O’M., H.W., D.M, X.W.; Investigation, J.Q., W-H.K, G.J.C, E.C, L.Z, A.M; Resources, Y-C.Y., Z.L.; Formal Analysis, R.C.R, S.B., Y-H.K; Writing-Original Draft, R.C.R and Y-H.K; Writing – Review & Editing, L.D.W, S.J.F, N.C., G.M.; Funding Acquisition, R.C.R, Y-H.K and G.M.

## STAR Online Methods

### Mice

To induce expression of CM, *Cbfb^+/56M^/Mx1-Cre* C57BL/6 mice (4-8 weeks old, including both females and males) were injected with 14 mg/kg poly (I:C) (InvivoGen, San Diego CA) every other day for 7 doses. Similarly treated, wild-type, *Cbfb^+/56M^ or Mx1-Cre* littermates were used as controls. As previously described(Cai et al., 2016), induction of CM expression results in expansion of pre-leukemic hematopoietic stem and progenitor cell (HSPC) populations in the bone marrow and subsequent development of overt leukemia characterized by >20% cKit+ leukmia blasts, increased white blood cell counts, and splenomegaly. Peripheral blood samples were taken via retro-orital venous sinus before induction and monthly thereafter for the duration of the experiment. All mice were maintained in an AAALAC-accredited animal facility, and all experimental procedures were performed in accordance with federal and state government guidelines and established institutional guidelines and protocols approved by the Institutional Animal Care and Use Committee at the Beckman Research Institute of City of Hope.

### Flow cytometry

PBMCs were stained in PBS with 0.5% BSA for 15 min on ice with fluorescently labeled antibodies for cell surface markers, including cKit, CD3, B220, CD11b, CD11c, CD71, Ter119. Phenotypic HSPC populations were defined as LSK (Lin^-^ckit^+^Sca1^+^); MP (Lin^-^ckit^+^Sca1^-^); Pre-GM (Lin^-^ ckit^+^Sca1^-^CD41^-^CD16/32^-/lo^CD105^-^CD150^-^); GMP (Lin^-^ckit^+^Sca1^-^CD41^-^CD16/32^+^CD150^-^); Pre-Meg/E (Lin^-^ckit^+^Sca1^-^CD41^-^CD16/32^-/lo^CD105^-^CD150^+^). All antibodies were purchased from BioLegend, BD Biosciences, or eBiosciences. Flow cytometry was performed using a 5-laser, 15-detector BD LSRII or LSRFortessa. Data analysis was performed using FlowJo (Tree Star, Ashland OR).

### RNA extraction, sequencing, and bioinformatics

Peripheral blood samples were accrued for all timepoints and allocated to randomized batches for RNA extraction. Total RNA was extracted from whole blood using the AllPrep DNA/RNA Mini Kit (Qiagen, Hilden, Germany); quality and quantity were estimated using a BioAnalyser (Agilent, Santa Clara, CA). Samples with a RIN > 4.0 were included. External RNA Controls Consortium (ERCC) Spike-In Control Mix (Ambion, Foster City, CA) was added to all samples per the manufacturer’s recommendations, although these were not used for downstream analyses. Samples were allocated to randomized batches for library preparation, such that samples from each timepoint were distributed evenly over all sequencing runs. Sequencing libraries were constructed using the KapaHyper with RiboErase (Kapa Biosystems, Wilmington, MA), loaded on to a cBot system for cluster generation, and sequenced on a Hiseq 2500 (Illumina) with single end 51-bp for mRNA-seq to a nominal depth of 40 million reads. To mitigate batch effects, samples were randomly assigned to one of eight flow cells such that each flow cell contained a sample from at least one mouse and one timepoint. Image processing and base calling were conducted using Illumina’s Real-Time Analysis pipeline.

Raw sequencing reads were processed with the nf-core RNASeq pipeline version 1.0 (Ewels et al., 2018). Briefly, trimmed reads were mapped using Spliced Transcripts Alignment to a Reference (STAR)(Dobin et al., 2013) to the GRCm38 reference amended with ERCC sequences and the human *MYH11* fusion gene sequence. Each library was subjected to extensive quality control, including estimation of library complexity, gene body coverage, and duplication rates, among other metrics detailed in the pipeline repository(Ewels et al., 2018). Reads were counted across genomic features using Subread featureCounts(Liao et al., 2014) and merged in to a matrix of counts per gene for each sampled timepoint. *CM* fusion transcript expression was measured by counting reads that spanned the *CM* fusion boundary. Surrogate variable analysis was used to check for confounding experimental effects(Leek et al., 2018). None were apparent (data not shown); however, library preparation batch was used as a covariate in differential expression analyses using the Bioconductor package edgeR(Robinson et al., 2009) as implemented in SARTools (Dobin et al., 2013).

### Supplemental methods and information

In this supplement we provide additional data to support our methods and results. In particular, we: show details of our mouse model of AML; demonstrate the robustness of our state-space construction to normalization methods and gene selection criteria; compare dimension reduction methods (PCA, Diffusion Mapping, t-SNE, MDS); show bootstrap cross-validation of our predictions; include detailed illustrations of individual state-transition trajectories and potential energy landscapes; compare hierarchical clustering with our state-space approach; and finally, provide full lists of differentially expressed genes and gene ontology (GO) term enrichment. Each of the concepts is illustrated with a figure.

### Uncertainty quantification and sensitivity analysis

To evaluate the sensitivity of our state-space construction to variations in sample frequency, number of reads per gene, and number of timepoints, each of these quantities was varied independently (Figure S4-6). Sample frequency was assessed by varying the gene inclusion criteria from 5 counts in at least 2 samples, to 5 counts in each of 5, 10, 30, 50, 70, 90, 110, 120, and all 132 samples, with 5 counts in 2 samples being the most permissive criteria, resulting in 21,482 genes, to 5 in 132 samples being the most restrictive criteria, resulting in 8,995 genes. The number of reads per gene was assessed by varying the log of the counts per million reads (cpm) log2(cpm) threshold in increments of 0.01, 0.05, 0.5, 5, 1, 3, 5, 7, 10, 15, and 20 for each of the sampling frequencies, resulting in 100 combinations. For a subset of combinations, the effect of normalization methods [e.g., trimmed means of M values (TMM), relative log expression (RLE), upper quartile (UQ), and RLE with ComBat regularization] on the state-space was examined. Sampling frequency during leukemia evolution was assessed by performing a leave- “x”-out cross-validation technique, in which x=70 samples were randomly identified and removed from the data set. The remaining 62 samples were used to predict the positions in the state-space for the 70 removed samples. This cross-validation procedure was performed 100 times and the absolute difference between the true state-space position and the predicted position was computed. The state-space was also constructed with and without the *CM* and *Kit*. There was no difference in state-space positions when the state-space was constructed without these genes, up to machine precision, 1×10^-14^ (Figure S9).

### Leukemia eigengene selection

Due to the large number of differentially expressed genes at the leukemic endpoint (c_3_), genes were filtered based on the angle subtended (*θ*) by the gene in the 2-dimensional state-space. The range of angles (*θ*) identified as being associated with leukemia evolution was defined as the minimal sector of a circle centered at (0,0) that included all leukemic samples as well as the mirror image of this sector along the x-axis of symmetry (Figure S19).

## SUPPLEMENTAL DATA FIGURES

**Figure S1.**
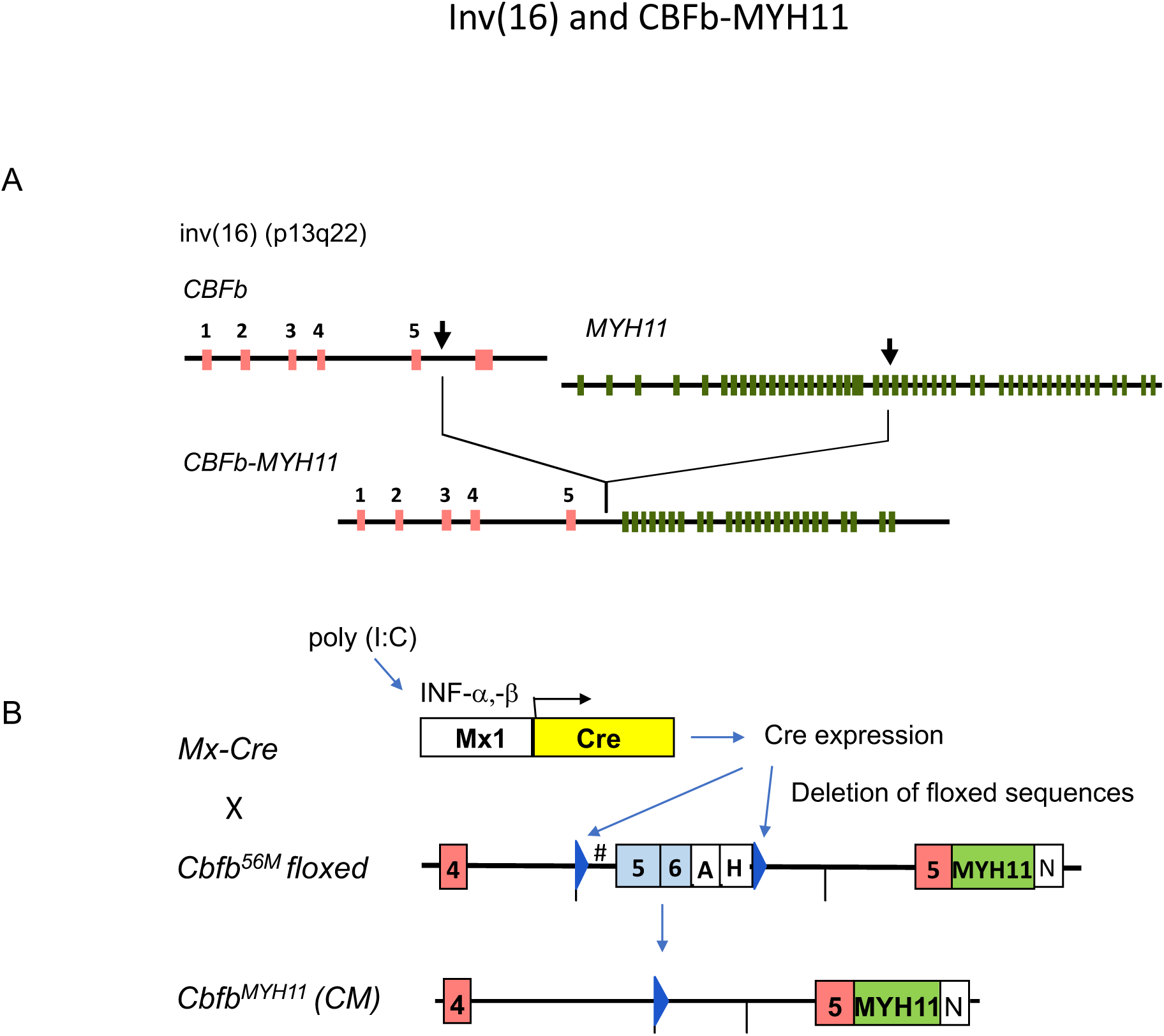
Inv(16)(p13q12) chromosomal rearrangement and the conditional knock-in for modeling AML induced by the *CBFb-MYH11* fusion gene. A) Illustration of chromosomal rearrangement inv(16) (p13q22), which fuses the *CBFb* gene with the *MYH11* gene, resulting in aberrant *CBFb-MYH11* fusion gene. B) Illustration of the conditional knock-in mouse model (*Cbfb^+/56M^/Mx1-Cre)* used to mimic inv(16) induced AML in this study. *Mx1-Cre* transgenic mice were bred with *Cbfb^+/56M^* floxed mice, carrying a knock-in of a loxP-floxed (blue triangle) *Cbfb* exon 5-6 cassette and human *MYH11* sequence fused with the exon 5 of mouse *Cbfb* and. Injection of a synthetic double-stranded RNA, poly (I:C), induces the interferon (INF) responsive promoter *Mx1-Cre* and thus expression of Cre recombinase, deleting the *Cbfb^56M^-*floxed sequences, and creating the *Cbfb-MYH11* fusion gene.

**Figure S2.**
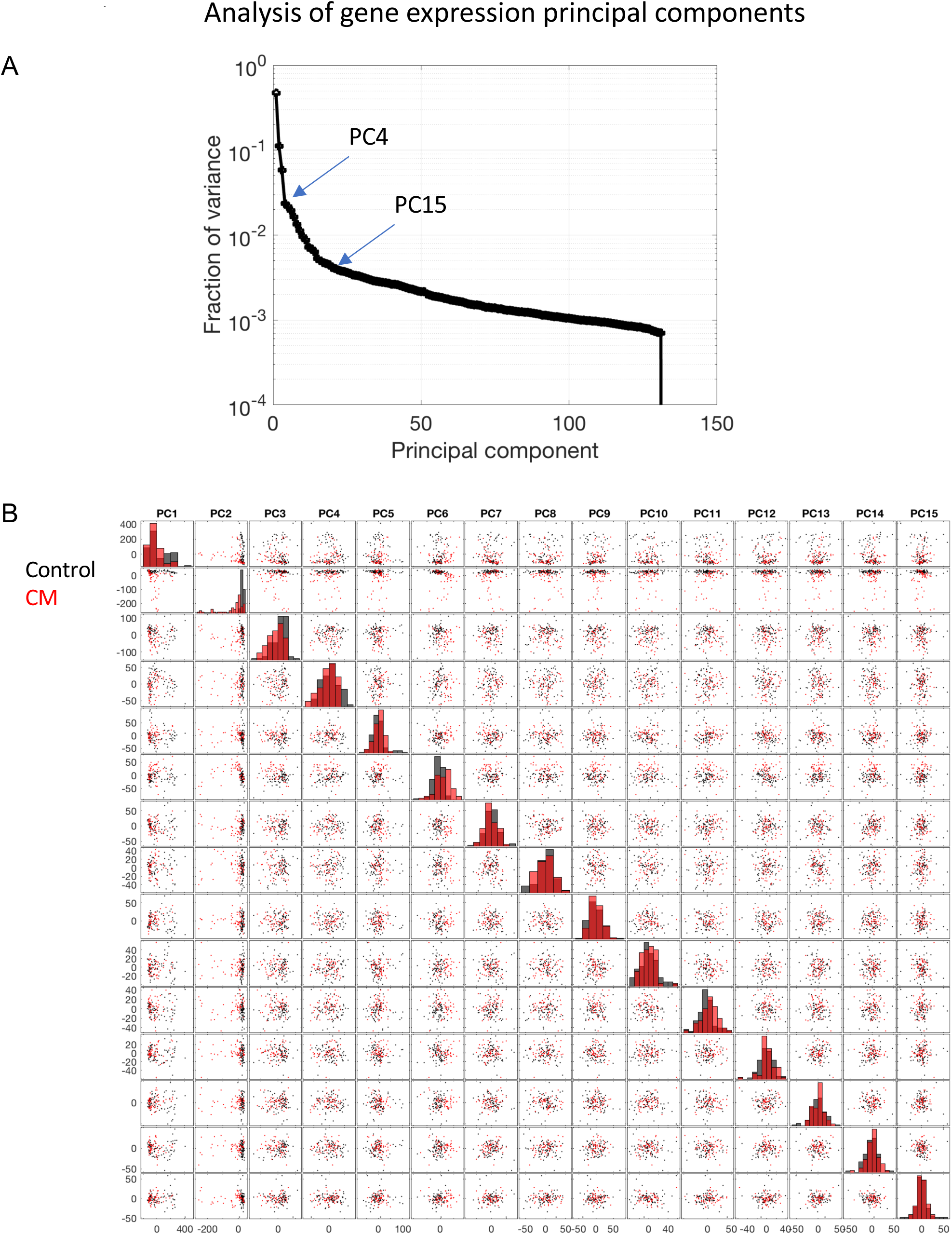
Analysis of gene expression principal components. A) Two elbows are identified in the principal component spectrum of the data matrix X; one at component 4 and a second at component 15. This indicates that information content is very low beyond PC15. B) A pairwise plot of each of the first 15 components. PC2 is the only component to segregate control (black) and CM (red) samples. This can be seen in the histograms along the diagonal. The second row demonstrates that any other component (PC1-PC15, exclusive of PC2) could be used as an orthogonal state to PC2 (leukemia) and that no other component encodes leukemia dynamics.

**Figure S3.**
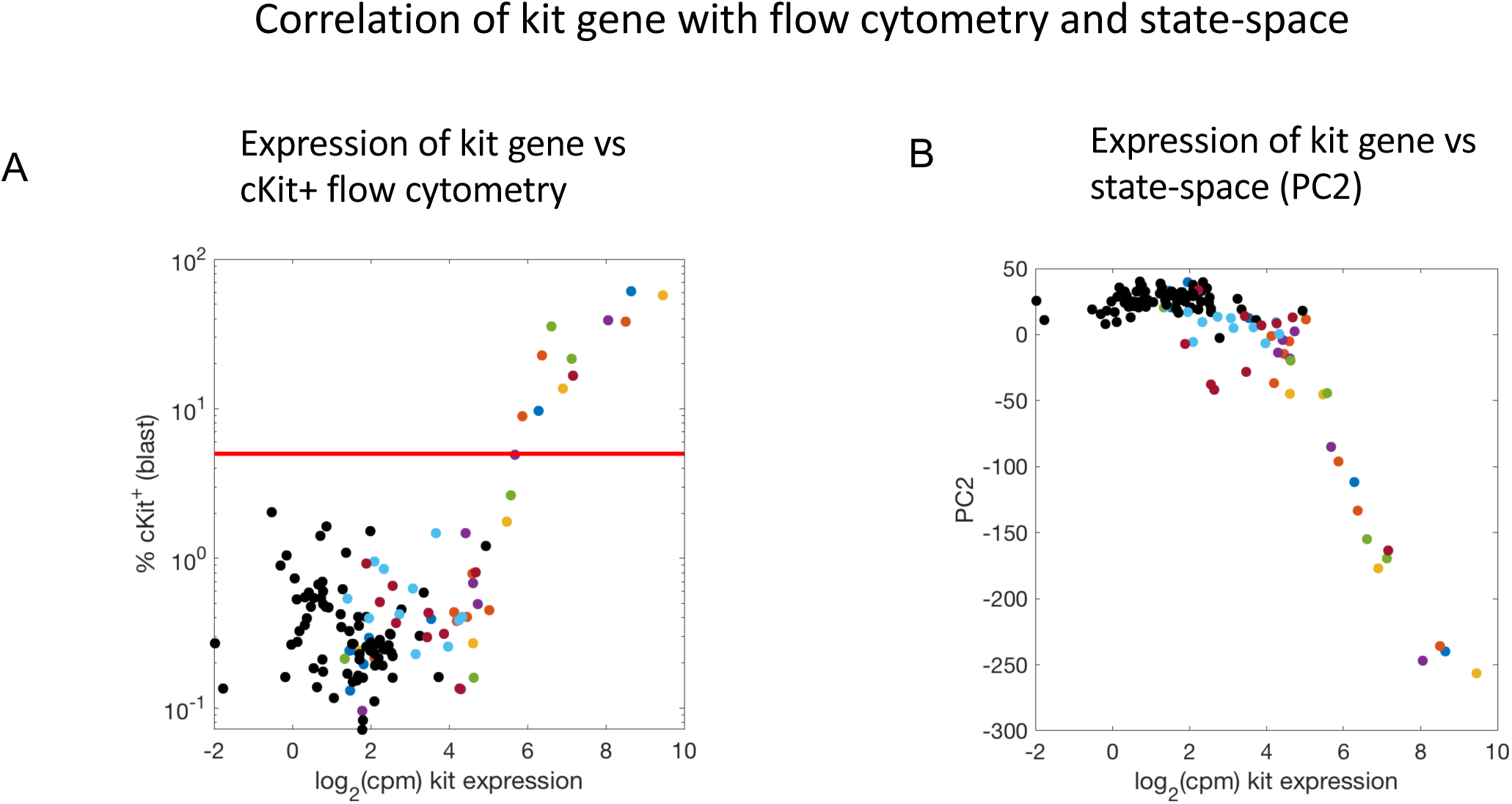
Correlation of kit gene with flow cytometry and state-space. A) expression of *kit* gene plotted against percent of cKit+ cells as determined with flow cytometry. Controls are shown in black and colors are coordinated for each CM mice. Red line indicates 5% cKit+ blast cells. Expression and FACS are correlated only for cKit+ > 5%. B) kit gene expression plotted against PC2. Colors same as in A).

**Figure S4.**
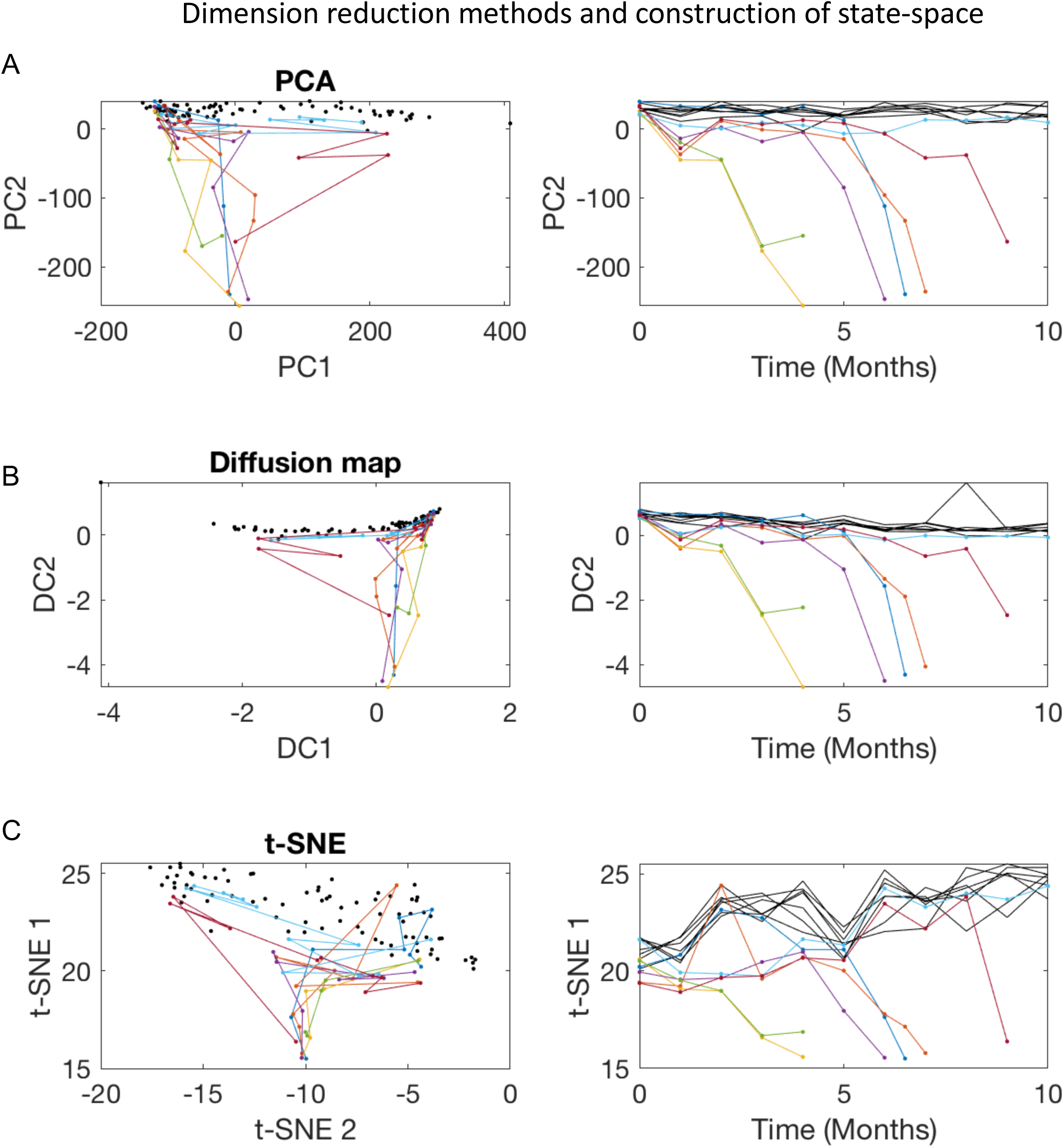
Dimension reduction methods and construction of state-space. Three methods of dimension reduction are compared to construct the leukemia state-space. The left column is the 2D space and the right column is the component which encodes leukemia plotted over time. Colors are matched in all plots. Each CM mice is assigned a color. Controls are shown in black. A) Principal component analysis (PCA). We use PCA in our analysis and results. B) Diffusion mapping. Trajectories are very similar to PCA, however, diffusion mapping involves 2 free parameters which must be tuned. C) t-SNE embedding. This approach does not clearly differentiate control and leukemia trajectories. PCA is preferable because: it has no free parameters (ex. diffusion maps); is not stochastic (ex. t-SNE); and is a linear map, which is more easily interpretable. Moreover, PCA can easily be used to map new data into an existing space; this is not easily achievable with other methods. Diffusion map and t-SNE implemented with MATLAB.

**Figure S5.**
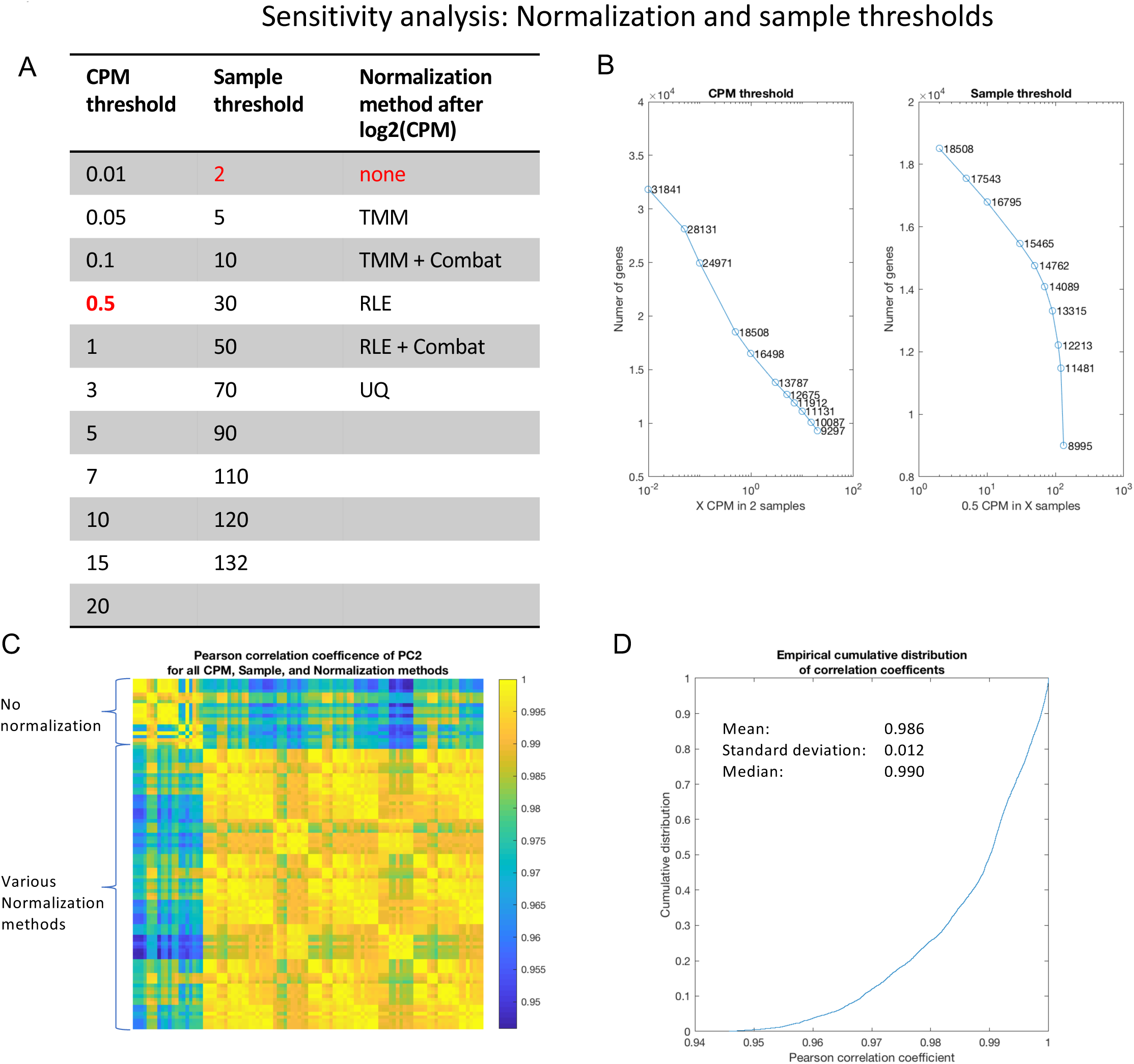
Sensitivity analysis: normalization and sample thresholds. We examined the impact of various normalization and sample thresholds on construction of the state-space with PCA. A) We varied the minimum counts per million reads (cpm) threshold for gene inclusion; the number of samples which a gene appears; and normalization methods. B) The number of genes included in the analysis is plotted against Left) various CPM thresholds for a fixed sample threshold and Right) the sample threshold for a fixed CPM threshold. C) All possible combinations of CPM, Sample, and Normalization methods are compared against each other. The colorbar indicates the Pearson correlation coefficient between each method, evaluated for PC2. The correlation is weakest when comparing state-spaces constructed with normalization to those without normalization (top rows; first columns), with correlation coefficients ranging from 0.95-0.995. When comparing state-spaces with normalization to each other, the correlation increases significantly. D) The empirical cumulative distribution of Pearson correlation coefficients for all combinations shown in C). The mean is 0.986 with standard deviation 0.012, indicating the state-space geometry is extremely robust to CPM, sample thresholds, and normalization methods. For our final analysis we used: 0.5 reads in at least 2 samples with no normalization method (red in A); although this analysis demonstrates that this choice has very little impact on the state-space geometry or results. Example trajectories are shown in Figure S5.

**Figure S6.**
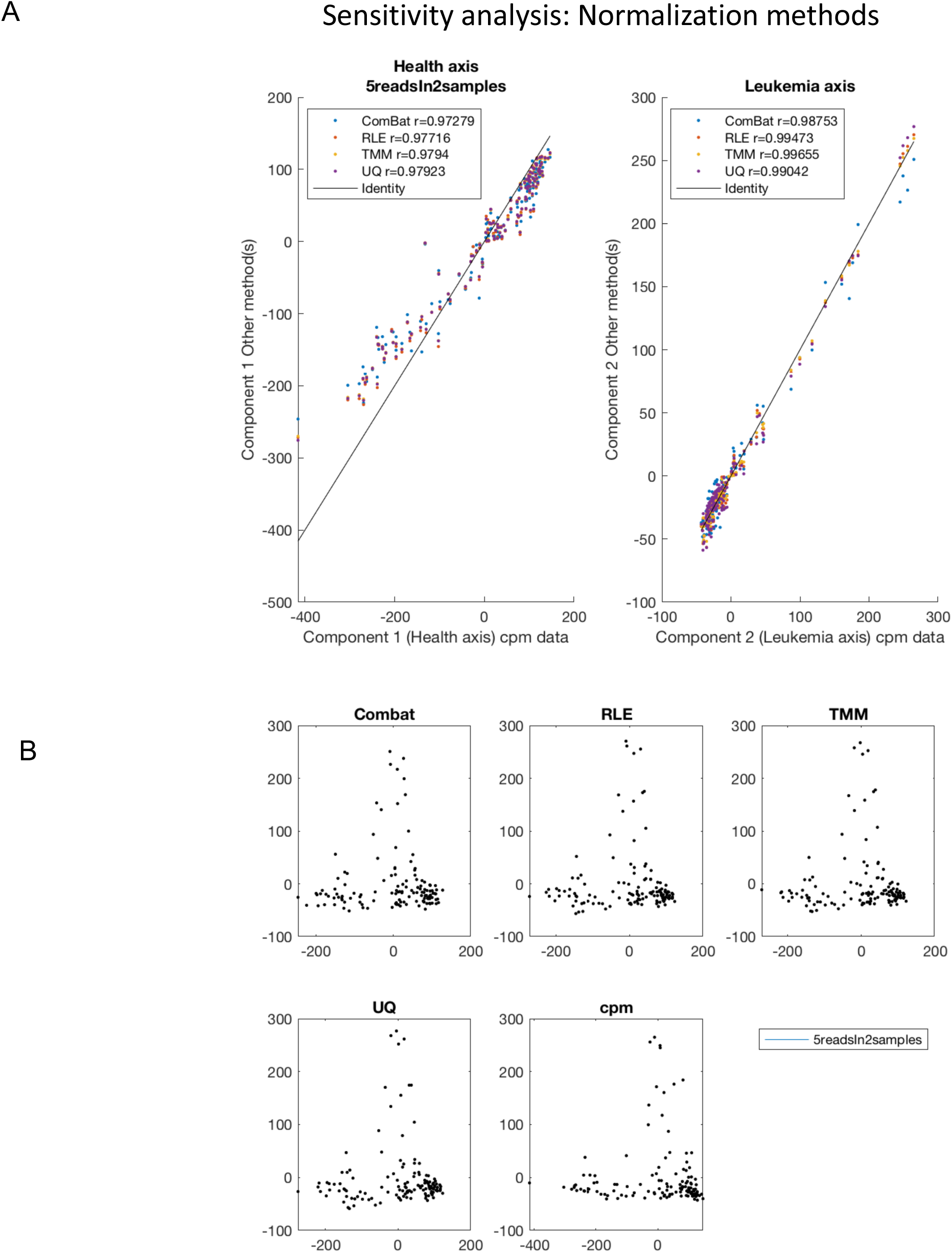
Sensitivity analysis: normalization methods. A) Correlation of 4 normalization methods and the reference (no normalization) is shown for PC1 (left) and PC2 (right). B) Example state-space geometries for 4 normalization methods (Combat; RLE; TMM; UQ) and no normalization (cpm) for a sample threshold of 5 reads in at least 2 samples. Note the state-space is inverted as compared to that shown in the main text. This is because principal components are uniquely determined up to a sign factor (+/- 1), which is arbitrarily chosen. The spaces are mathematically equivalent.

**Figure S7.**
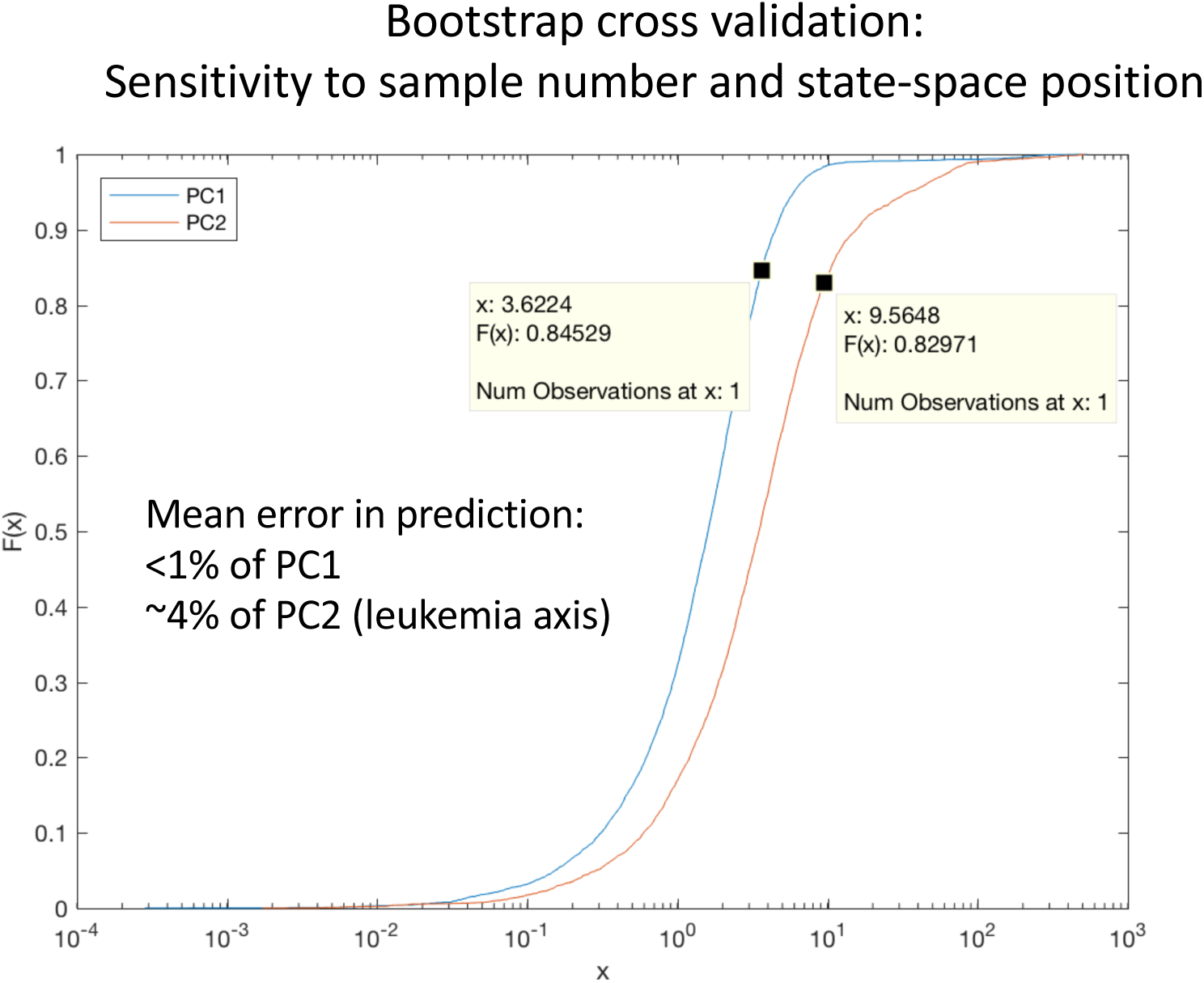
Bootstrap cross-validation. We performed bootstrap cross-validation with the 5 reads in at least 2 samples threshold and no normalization dataset with the following procedure: 1) randomly remove 70 of the 132 total samples; 2) build state-space with remaining 62 samples; 3) predict locations of 70 removed samples by mapping them into the space by multiplying by the conjugate transpose of the eigengene matrix V; 4) evaluate the difference between the actual and predicted locations in the state-space; 5) repeat the process 100 times. The empirical cumulative distribution of the results are shown. The mean error in prediction for PC1 is < 1% and ∼4% for PC2. Note the x-axis is log scaled. Data markers are placed at the arithmetic mean. The median value (x such that F(x)=0.5) is significantly smaller. This analysis demonstrates that the state-space construction is not sensitive to the inclusion of specific data points in the leukemia or control trajectories.

**Figure S8.**
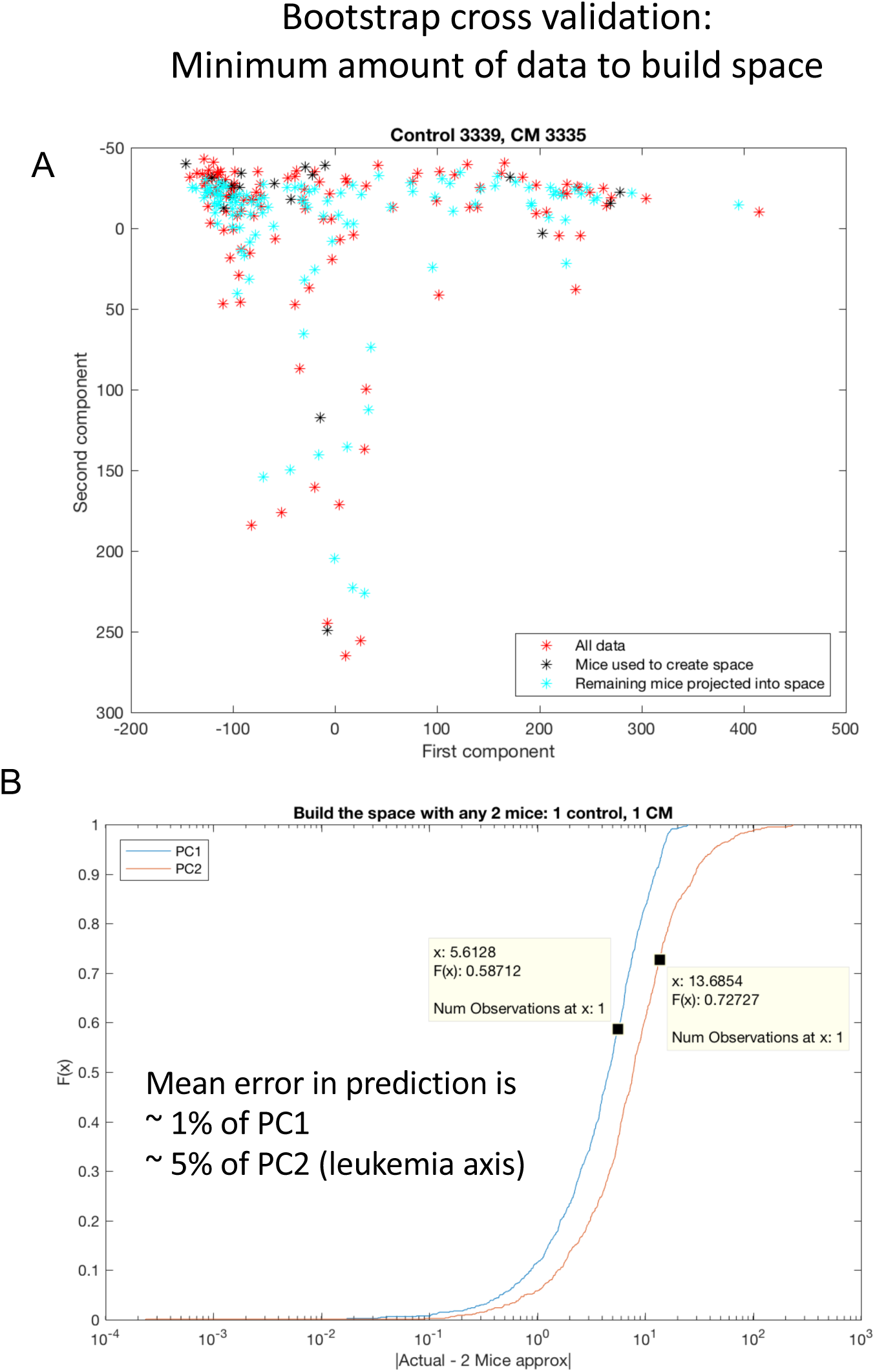
Bootstrap cross-validation. We performed an additional bootstrap cross-validation with the 5 reads in at least 2 samples threshold and no normalization dataset to determine the minimum amount of data required to construct the state-space with the following procedure, shown in A): Randomly choose 1 control and 1 CM mice 2) Build space with 2 mice (*) 3) Predict locations of all other samples (*) 4) Evaluate difference (* - *) 5) Repeat for all combinations of control and CM mice selections. B) The empirical cumulative distribution of the results are shown. The mean error in prediction for PC1 is ∼ 1% and ∼5% for PC2. Note the x-axis is log scaled. Data markers are placed at the arithmetic mean. The median value (x such that F(x)=0.5) is significantly smaller. This analysis demonstrates that the state-space may be constructed with temporal data obtained from as few as 2 mice: 1 control and 1 CM to construct orthogonal states. This is sufficient to predict the positions and dynamics of data not used to construct the space.

**Figure S9.**
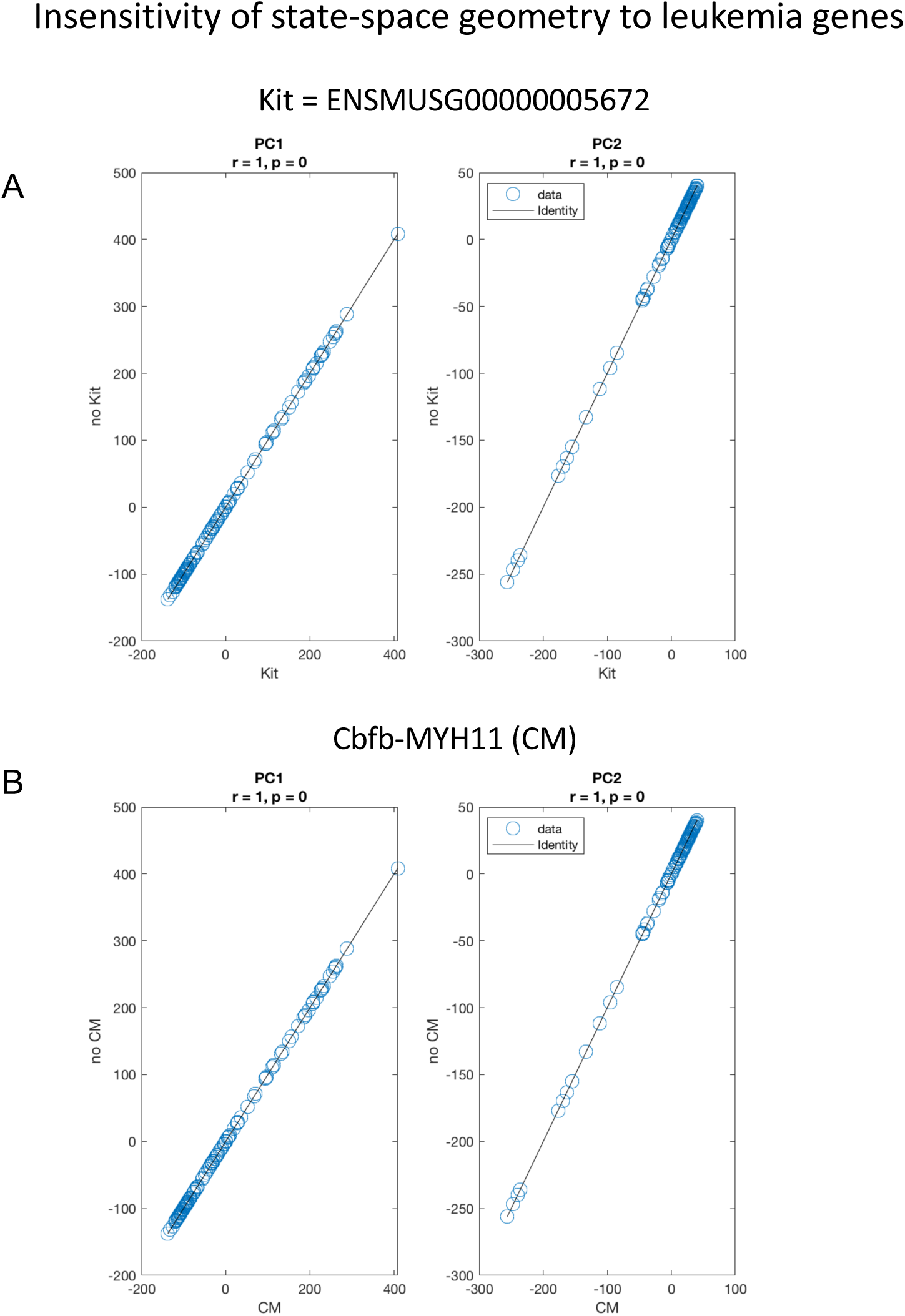
Insensitivity of state-space geometry to known leukemia genes. We asked the question to what extent the state-space geometry is determined by known leukemia driver genes *Kit* and *Cbfb-MYH11 (CM)*. To answer this question we removed each of these genes from the data set and compared the state space geometry (PC1, PC2) with (x-axis) and without (y-axis) these genes. A) With and without *Kit* B) with and without *Cbfb-MYH11*. Differences are so small the correlation coefficient is rounded to unity. No differences are detected. The line of identity is plotted for reference.

**Figure S10.**
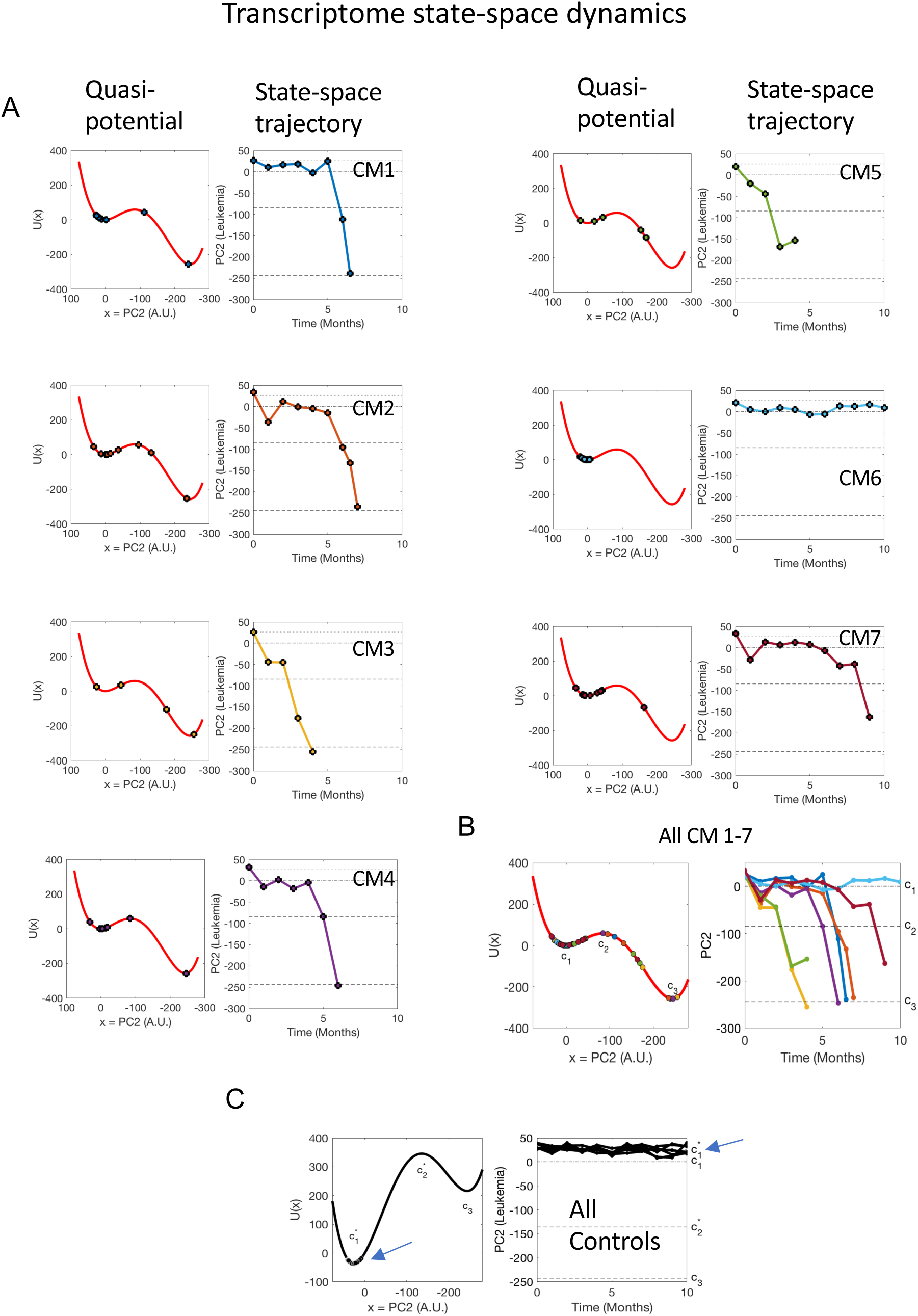
Transcriptome state-space dynamics. Left column) The quasi-potential energy *U(x)*, with each point *x_i_* evaluated on the potential *U(x_i_)*. Right column) PC2 state-space trajectory. A) Trajectories and potential energy values for each of the 7 CM mice, shown individually. Critical points c_1_, c_2_, c_3_ are shown as dashed lines in state-space trajectories. B) All 7 CM mice plotted together. Colors are coordinated with A). C) All control mice trajectories are in the local minimum at c_1_*.

**Figure S11.**
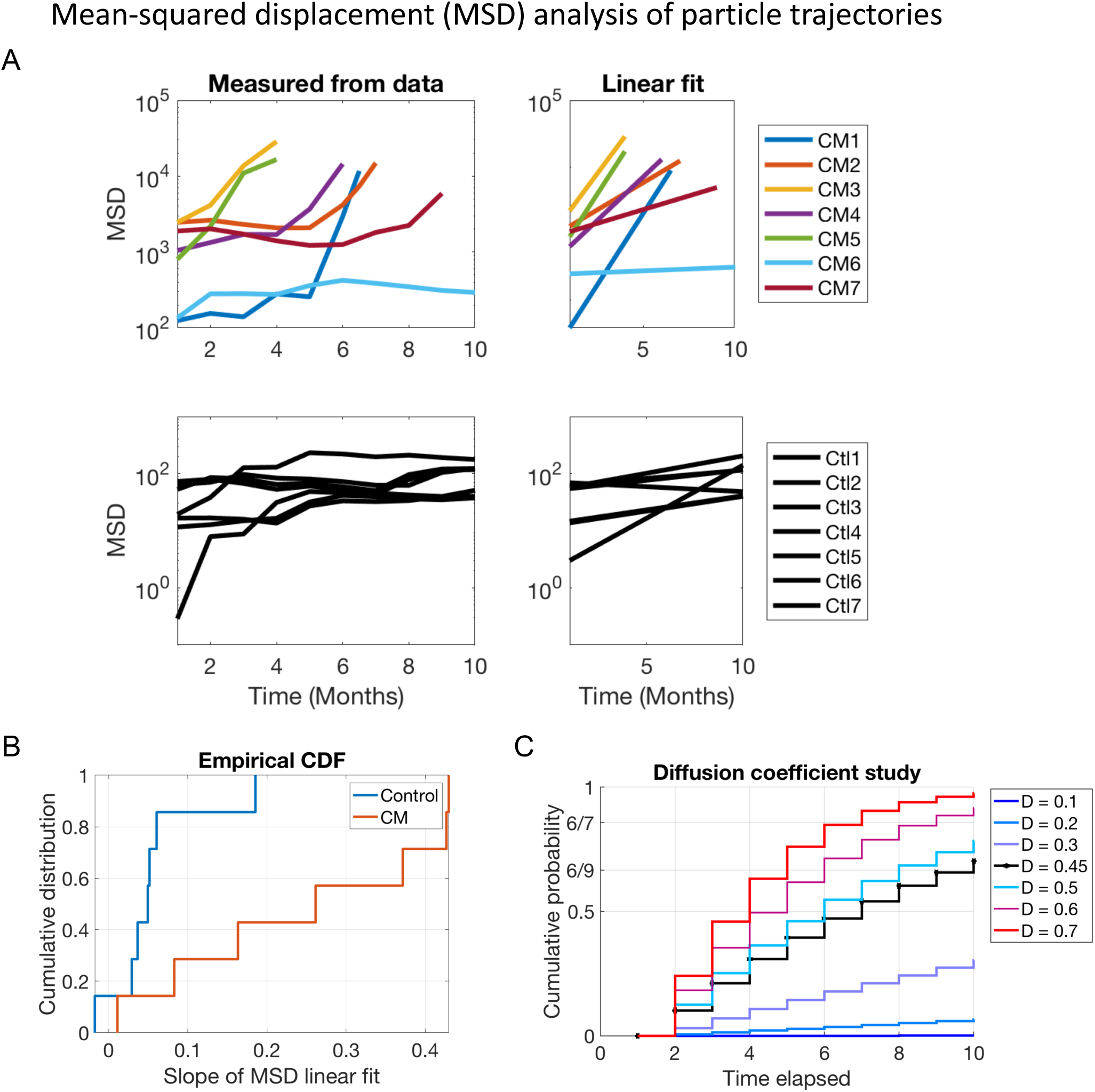
Mean-squared displacement (MSD) analysis of particle trajectories. A) MSD analysis was performed to estimate the diffusion coefficient of particle trajectories for the training cohort. The top row is CM mice and the bottom row is controls. The left column is MSD calculated from PC2 trajectories from the training cohort and the right column is a linear fit for each mouse trajectory. The slope of the linear regression gives an estimate for the parameter *D* = *β*^−1^. The average slope of the linear regression was used as the diffusion coefficient in the stochastic simulations for CM and control mice respectively (Figure 5A) and the Fokker-Planck (FP) equation (Figure 5B). B) The empirical cumulative distribution function of slopes of linear fits shown in A). Control trajectories have smaller slopes than CM, reflecting restricted movement (diffusion). The diffusion coefficient for CM mice was estimated as the mean value of the slopes of the linear fits to the MSD curves for CM mice, *D* = 0.25 (x^2^/time). (C) The FP is solved for a range of diffusion coefficients (D=[0.1-0.7]) derived from the MSD analysis (x-axis of B), and the cumulative probability of state transition to leukemia computed. The diffusion coefficient of D = 0.45 (x^2^/time) was used based on the training cohort and applied to predict the validation cohorts with survival analyses where survival represents the time to develop leukemia: for mice this is defined as >20% blast in peripheral blood, for simulation, this is defined as the integral probability mass past the critical point c_2_ (Figures 5,7).

**Figure S12.**
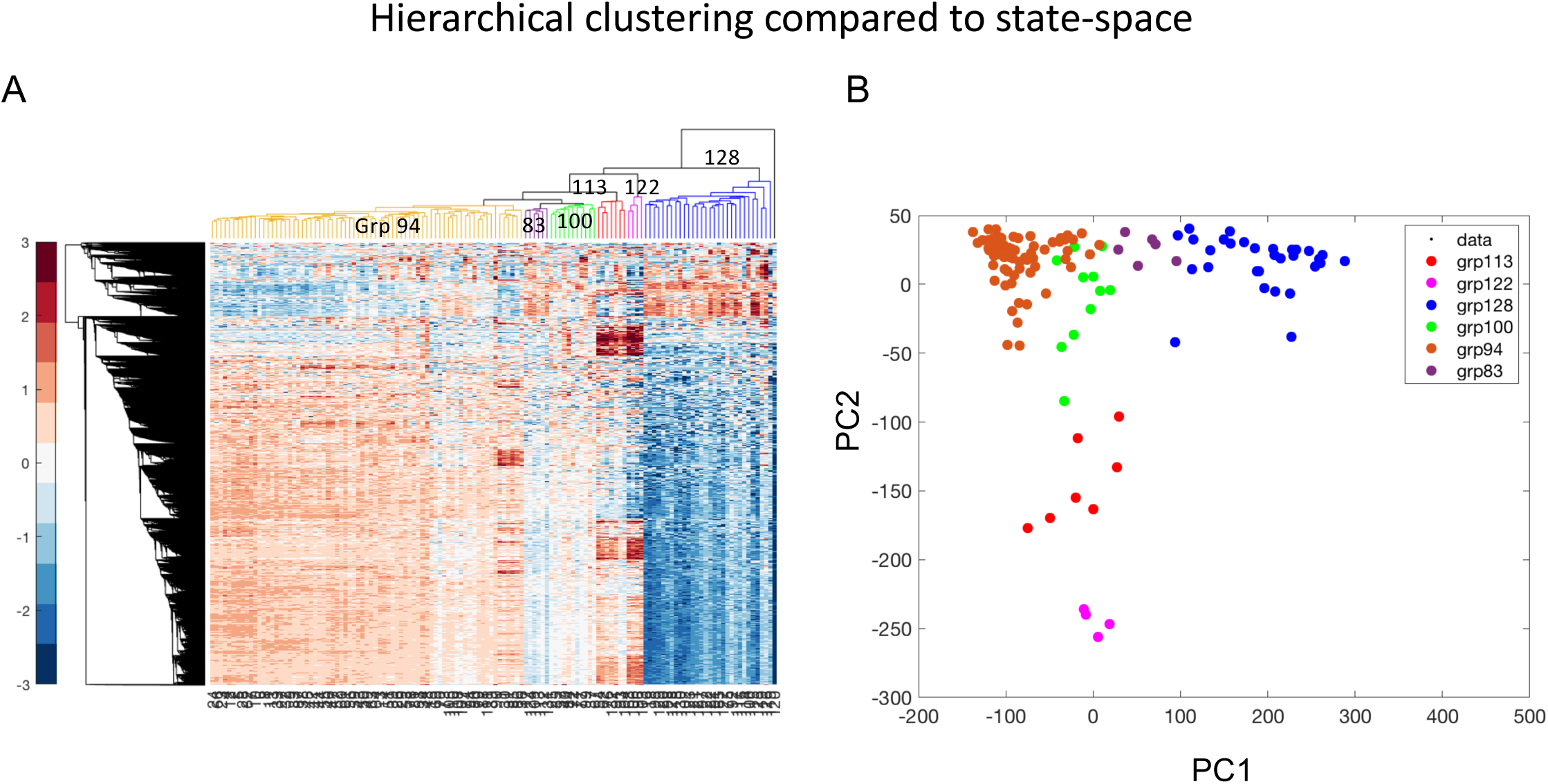
Hierarchical clustering compared to state-space. A) We performed hierarchical clustering on both the rows and columns of the gene expression data matrix X. B) We mapped the clusters into the 2D PCA state-space. Clustering correctly identifies the final leukemic samples (grp122), but fails to differentiate controls from CM mice, and fails to partition the samples temporally. This is seen by the predominant clustering in the state-space along PC1 rather than PC2. In particular, groups 113, 100, and 128 include control and CM mice and span a range of PC2 from [-50, 50].

**Figure S13.**
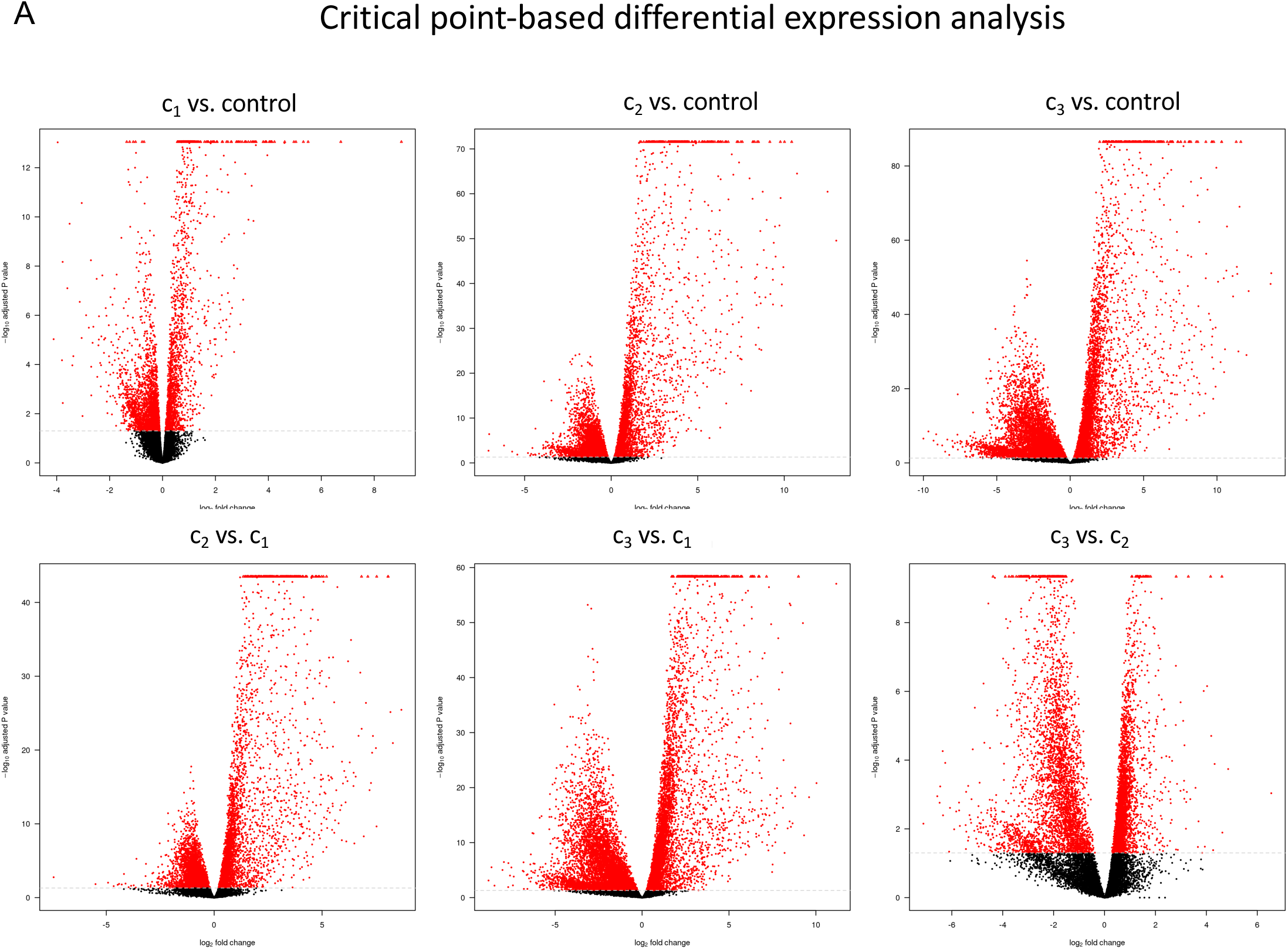
Critical point-based differential expression analysis. State-transition critical points were used to partition the samples for differential expression analysis. Samples were uniquely identified with a critical point by selecting the point with the minimum distance between the sample and the critical points (c*_1_,c_1_,c_2_,c_3_). A) Log2 fold change is plotted against –Log10 of the multiple test corrected p-value (q-value) for each comparison, commonly known as a volcano plot. Red indicates statistically significant differential expression, with significance level q < 0.05. Note that over 70% of sequenced genes are differentially expressed at the leukemic endpoint (c_3_) as compared to control.

**Figure S14.**
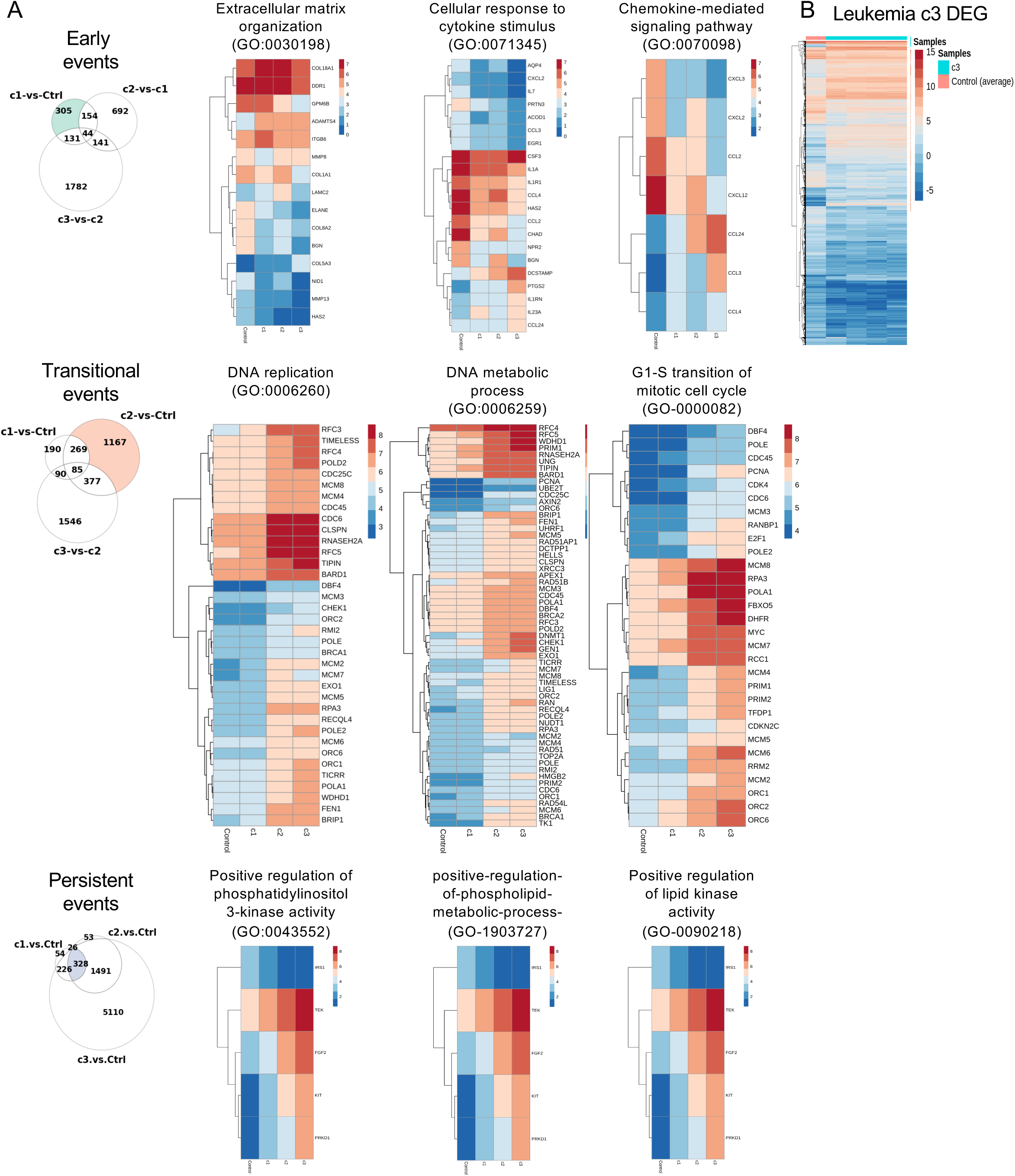
State-transition based analysis of genes and pathways in leukemia progression. A) Early, transition, and persistent events in leukemia progression are defined relative to critical points. Gene expression heatmap of each DEG in the top 3 GO term ranked by q-value. B) Heatmap for all leukemia c3 DEG compared to the average of control samples.

**Figure S15.**
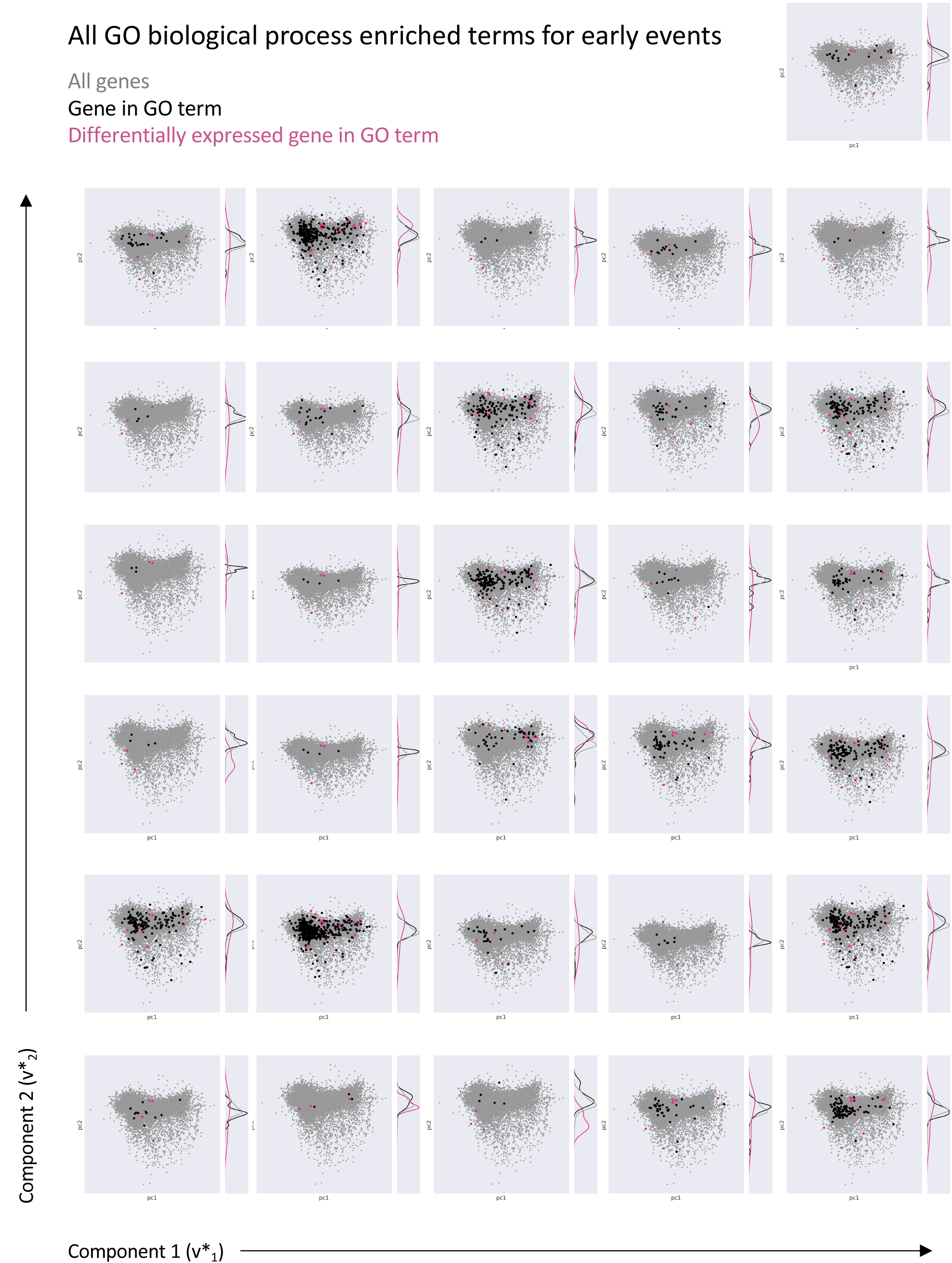
Eigengene representation of GO terms for early events. Each GO term is represented in the 2D eigengene space as in Figure 6D. Grey dots are all genes. Black dots are genes in the GO term. Pink dots are genes in the GO term which are differentially expressed in the early events samples. Each 2D eigenspace has a kernel density plot along component 2 to illustrate the locations in the eigengene space of the genes in the GO term. South is associated with leukemia.

**Figure S16.**
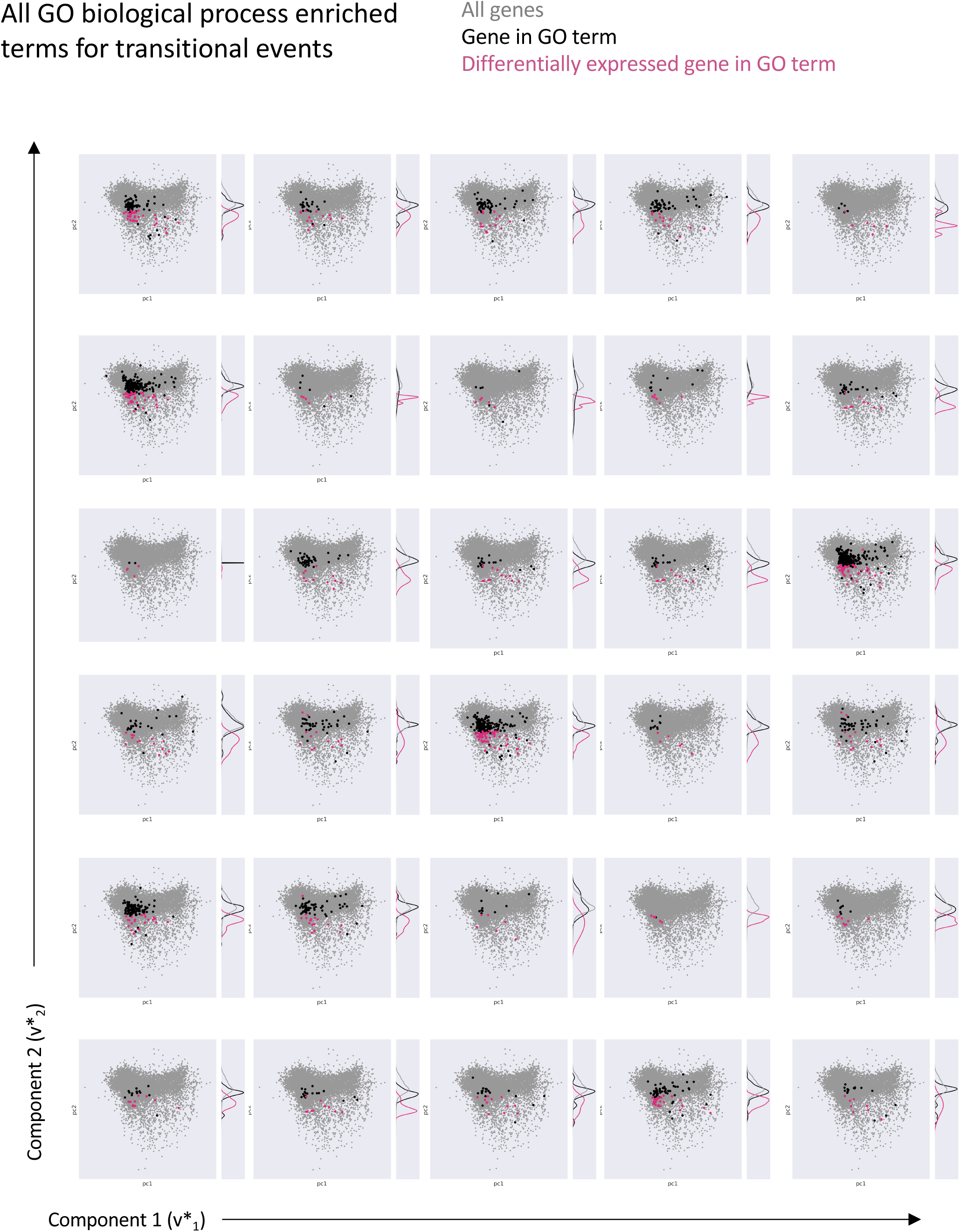
Eigengene representation of GO terms for transitional events. Each GO term is represented in the 2D eigengene space as in Figure 6D. Grey dots are all genes. Black dots are genes in the GO term. Pink dots are genes in the GO term which are differentially expressed in the transitional events samples. Each 2D eigenspace has a kernel density plot along component 2 to illustrate the locations in the eigengene space of the genes in the GO term. South is associated with leukemia.

**Figure S17.**
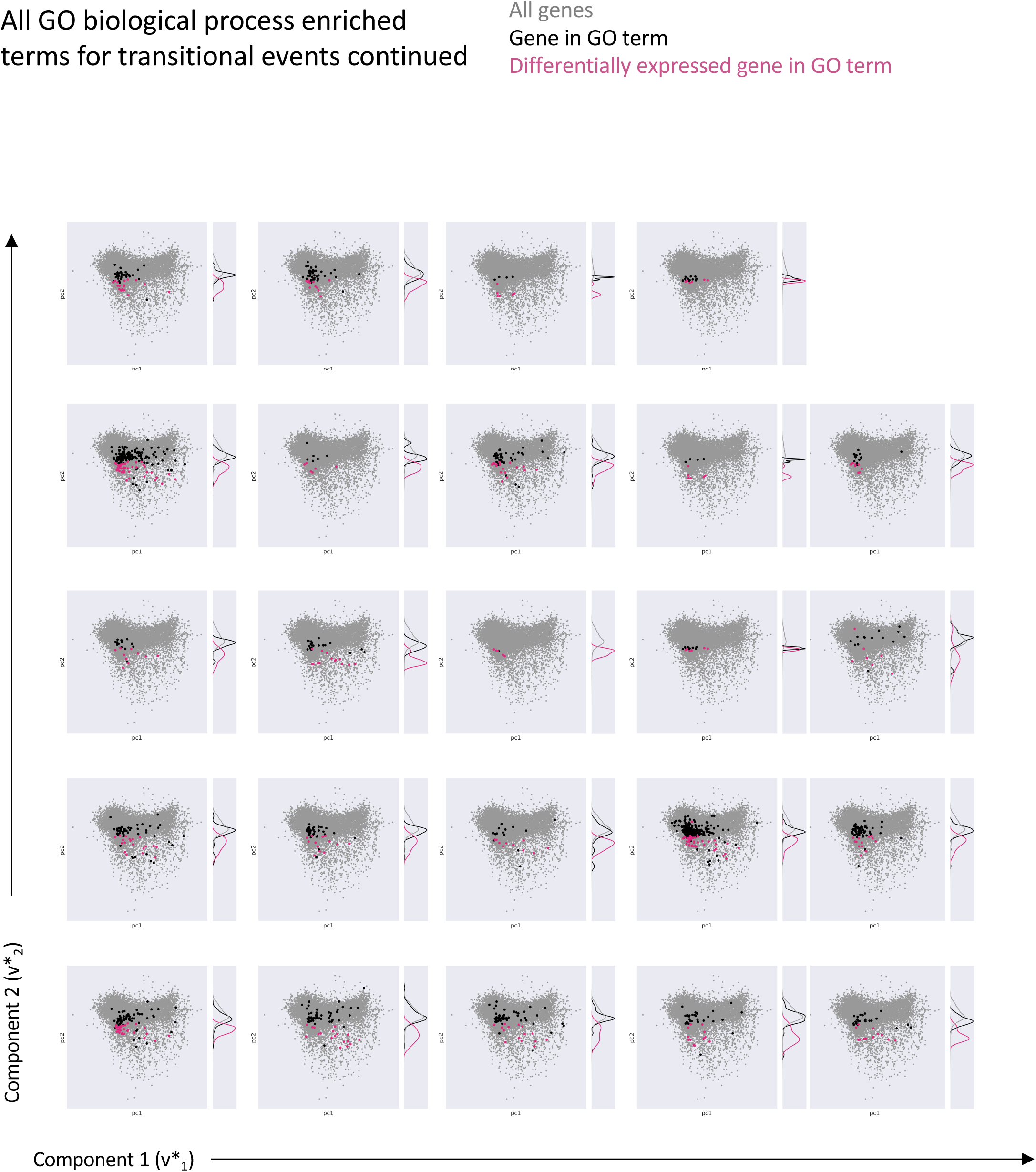
Eigengene representation of GO terms for transitional events – continued from S16.

**Figure S18.**
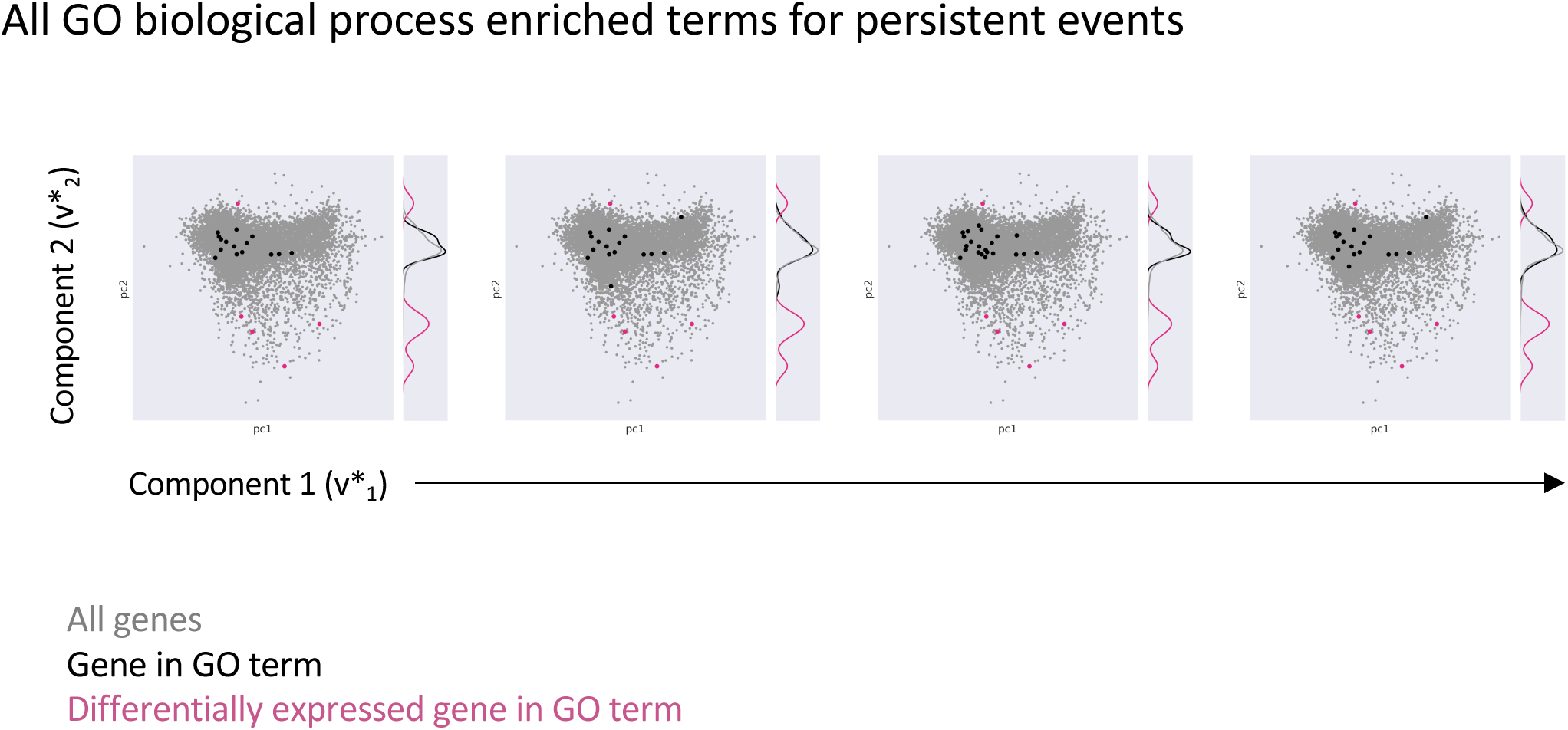
Eigengene representation of GO terms for persistent events. Each GO term is represented in the 2D eigengene space as in Figure 6D. Grey dots are all genes. Black dots are genes in the GO term. Pink dots are genes in the GO term which are differentially expressed in the persistent events samples. Each 2D eigenspace has a kernel density plot along component 2 to illustrate the locations in the eigengene space of the genes in the GO term. South is associated with leukemia.

**Figure S19.**
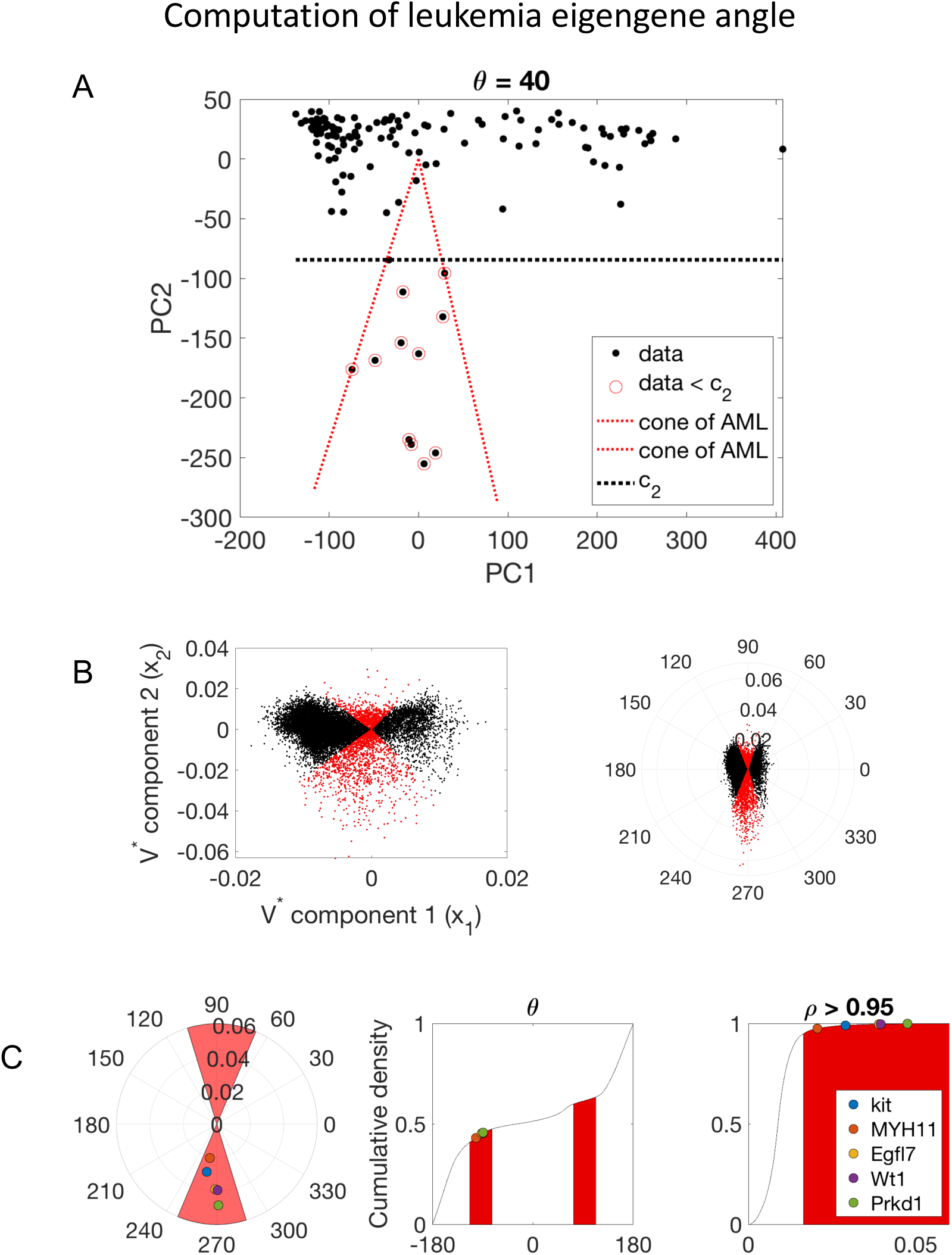
Computation of leukemia eigengene angle. (A) The geometric interpretation of the leukemia state-space (PC1,PC2) is used to identify a range of angles which are most strongly associated with state-transition to leukemia. The angle is calculated as the angle subtended by the minimum cone which includes all data points less than the critical transition point c_2_, centered at the center of mass of the data located at (0,0). The angle is approximately 40’. The cone is symmetric about the x-axis. (B) (Left) The first two columns of the eigengene matrix (V* see Figure 2A,F) can be represented in polar coordinates (ρ,θ,) (Right) to make more clear the range of angles. The leukemia eigengenes are genes which are in the leukemia cone (red). (C) Genes (*kit, MYH11, Egfl7, Wt1, and Prkd1*) shown in the leukemic cone in a polar plot (red area). Leukemia eigenegenes can also be represented as genes under the cumulative density curve for angle (ρ) or radius (θ,). There are 1,877 leukemia eigenegenes which fall into the leukemia cone, out of 18,508 genes sequenced, representing approximately 10% of sequenced genes. This novel analysis combines the geometry of the state-space with the eigengene space to select genes which most strongly associate with leukemia state-transition.

**Figure S20.**
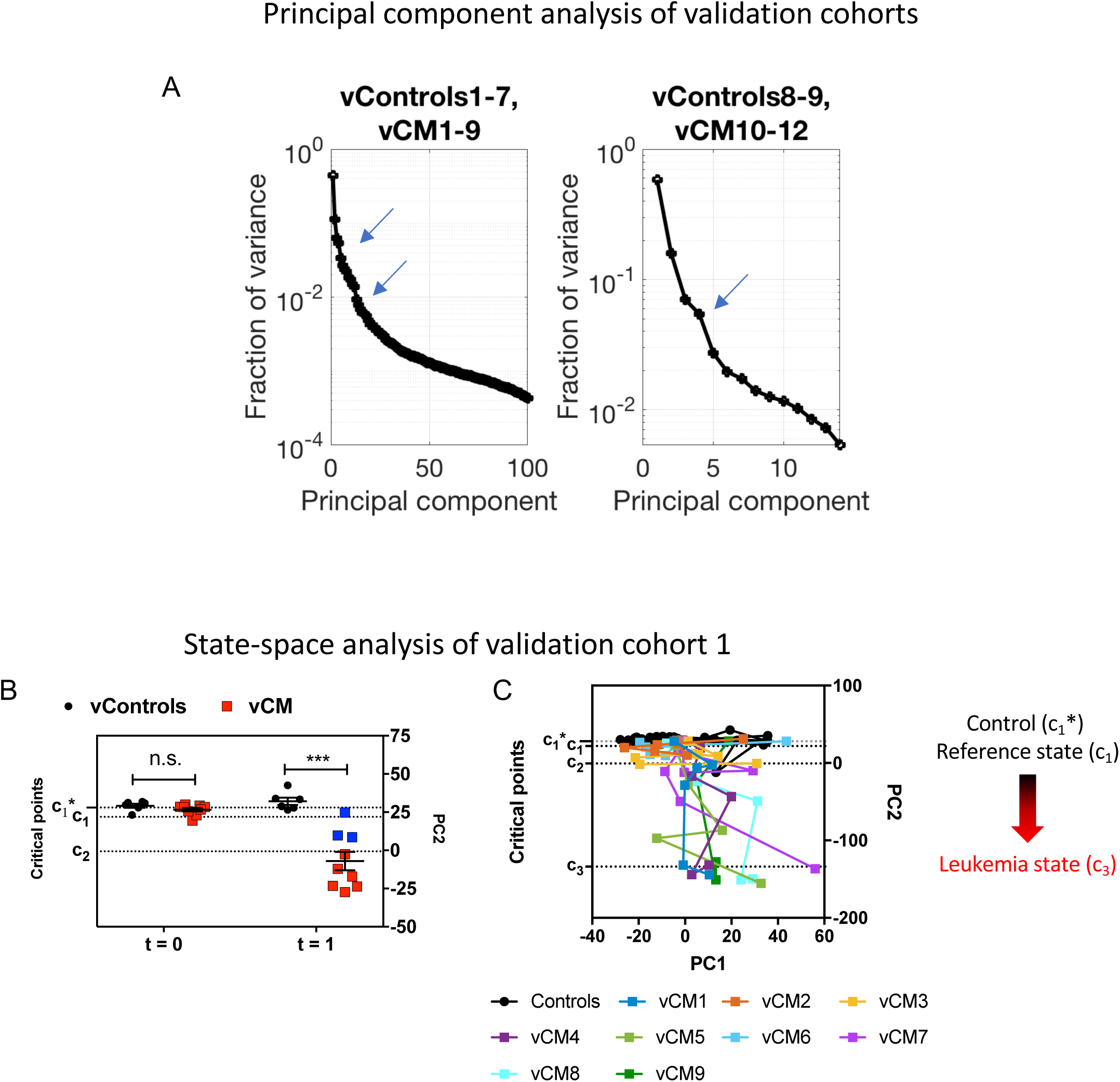
Principal component and state-space analysis of validation cohorts. (A) We performed component analysis of the validation cohort data matrices. The spectrum of the data matrices for each validation cohort is shown with the PCA “elbow(s)” indicated with arrows. The validation data sets show similar spectral distributions as is seen in the training dataset, namely that a majority of the variance is encoded within the first 4 PCs. (B) The validation cohort 1 (vControls1-7, vCM1-9) demonstrates the same movement in the state-space at t=1 as compared to t=0, reflecting the same CM oncogenic perturbation as seen in the training cohort (Figure 4D). The 3 vCM mice that do not develop leukemia during the experiment do not experience as significant drop in the state-space position (vCM not sick; blue). (C) Two-dimensional state-space trajectories with critical points estimated as in the training data.

**Figure S21.**
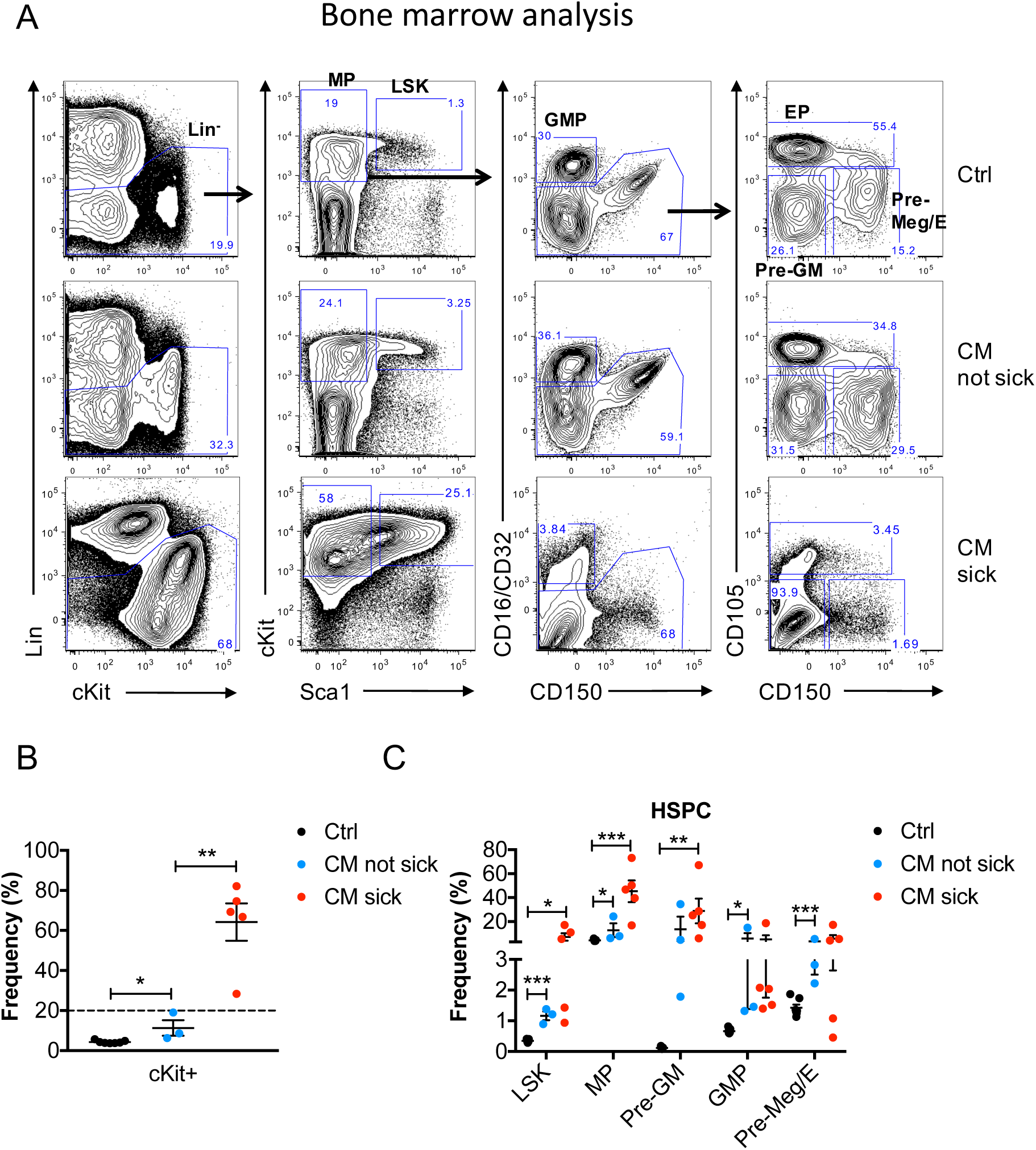
Phenotypic analysis of bone marrow (BM) at experimental end point (validation cohort). (A) Representative flow cytometry plots showing the gating strategy of various phenotypic progenitor populations including lineage-negative (Lin^-^), LSK (Lin^-^ckit^+^Sca1^+^); myeloid progenitor (MP; Lin^-^ckit^+^Sca1^-^); pre-granulocyte-macrophage (Pre-GM; Lin^-^cKit^+^Sca1^-^CD16/32^-/lo^CD150^-^CD105^-^), granulocyte-macrophage progenitors (GMP; Lin^-^cKit^+^Sca1^-^CD16/32^hi^CD150^-^), pre-megakaryocyte/erythrocyte (Pre-Meg/E: Lin^-^ cKit^+^Sca1^-^CD16/32^-/lo^CD150^+^CD105^-^), and erythroid progenitors (EPs: Lin^-^cKit^+^Sca1^-^CD16/32^-/lo^CD105^hi^) in the BM of moribund CM mice (CM sick), CM mice that had not developed leukemia at 6.5 months (CM not sick) compared to Ctrl mice analyzed at the experiment end point. The frequency within total BM is shown for Lin^-^ (first plot from left), and the frequency in the respective parent gate is shown for the other populations. (B) The frequency of cKit^+^ cells in the BM of moribund CM sick (n=6), CM not sick (n=3) compared to control (Ctrl) mice (n=7) at end point. Leukemia features expansion of immature cKit^+^ blasts (>20%, indicated by dished line) in both PB and BM. (C) The frequency of phenotypic progenitor populations in the CM sick (n=5), CM not sick (n=3) compared to Ctrl mice (n=7) BM as defined above. Increase of stem and progenitor cell populations (including LSK, MP, Pre-GM, GMP, Pre-Meg/E) were observed in all 3 CM mice that had not developed leukemia at 6.5 months. Mean ± SEM is shown. *: *p*<0.05; **: *p*< 0.01; ***: *p*<0.001.

Table S1. Differentially expressed genes for *c*_1_ vs 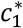

Table S2. Differentially expressed genes for *c*_2_ vs 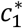

Table S3. Differentially expressed genes for *c*_3_ vs 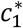

Table S4. Differentially expressed genes for *c*_2_ vs *c*_1_

Table S5. Differentially expressed genes for *c*_3_ vs *c*_1_

Table S6. Differentially expressed genes for *c*_3_ vs *c*_2_

**Table S7.** GO terms enriched for early events.

| Term | Overlap | P-value | Adjusted P-value | Genes |
| --- | --- | --- | --- | --- |
| extracellular matrix organization (GO:0030198) | 15/230 | 2.6866E-06 | 0.004231418 | DDR1;COL18A1;BGN;LAMC2;MMP8;NID1;ADAMTS4;COL1A1;MMP13;COL5A3;COL8A2;HAS2;ITGB6;GPM6B;ELANE |
| cellular response to cytokine stimulus (GO:0071345) | 21/457 | 7.9471E-06 | 0.006258322 | CCL24;EGR1;IL1RN;CSF3;IL1R1;NPR2;BGN;ACOD1;AQP4;PTGS2;CXCL2;IL1A;DCSTAMP;IL7;IL23A;CCL4;CHAD;CCL3;CCL2;HAS2;PRTN3 |
| chemokine-mediated signaling pathway (GO:0070098) | 7/53 | 1.5133E-05 | 0.006332687 | CCL24;CXCL12;CCL4;CCL3;CCL2;CXCL3;CXCL2 |
| positive regulation of calcium ion import (GO:0090280) | 4/11 | 1.6083E-05 | 0.006332687 | PDGFRB;CXCL12;CCL3;CCL2 |
| skin development (GO:0043588) | 8/95 | 0.0001028 | 0.02251213 | COL1A1;SPRR3;LCE3A;WNT10A;SPRR2G;CDSN;COL5A3;SPRR2D |
| positive regulation of cell motility (GO:2000147) | 11/180 | 0.0001043 | 0.02251213 | COL1A1;PDGFRB;CCL24;BMP2;DAB2;CXCL12;CCL3;SEMA3G;HAS2;LAMC2;IGF1 |
| cellular response to interleukin-1 (GO:0071347) | 10/151 | 0.00011073 | 0.02251213 | IL1A;CCL24;EGR1;IL1RN;IL1R1;CCL4;CCL3;CCL2;HAS2;ACOD1 |
| positive regulation of calcium ion transport (GO:0051928) | 5/34 | 0.00015451 | 0.02251213 | PDGFRB;CXCL12;CCL4;CCL3;CCL2 |
| positive regulation of cell migration (GO:0030335) | 12/222 | 0.00016356 | 0.02251213 | COL1A1;PDGFRB;CCL24;BMP2;DAB2;CXCL12;CCL3;SEMA3G;HAS2;LAMC2;IGF1;FGF10 |
| response to interleukin-1 (GO:0070555) | 7/77 | 0.00017285 | 0.02251213 | CCL24;IL1R1;CCL4;CCL3;CCL2;HAS2;ACOD1 |
| positive regulation of apoptotic cell clearance (GO:2000427) | 3/8 | 0.00018581 | 0.02251213 | C4B;C3;CCL2 |
| regulation of natural killer cell chemotaxis (GO:2000501) | 3/8 | 0.00018581 | 0.02251213 | CCL4;CCL3;CCL2 |
| smooth muscle cell migration (GO:0014909) | 3/8 | 0.00018581 | 0.02251213 | DDR1;PDGFRB;PLAT |
| positive regulation of smooth muscle cell proliferation (GO:0048661) | 5/38 | 0.00026518 | 0.02647997 | PDGFRB;NR4A3;HTR1B;IGF1;ELANE |
| neutrophil migration (GO:1990266) | 6/59 | 0.00027361 | 0.02647997 | CCL24;CCL4;CCL3;CCL2;PRTN3;CXCL3 |
| regulation of apoptotic cell clearance (GO:2000425) | 3/9 | 0.00027558 | 0.02647997 | C4B;C3;CCL2 |
| positive regulation of ERK1 and ERK2 cascade (GO:0070374) | 11/202 | 0.00028582 | 0.02647997 | DDR1;PDGFRB;IL1A;CCL24;BMP2;CCL4;CCL3;CCL2;TREM2;MERTK;FGF10 |
| regulation of calcium ion import (GO:0090279) | 4/23 | 0.00037349 | 0.032680385 | PDGFRB;CXCL12;CCL3;CCL2 |
| regulation of ERK1 and ERK2 cascade (GO:0070372) | 12/248 | 0.00044972 | 0.034253408 | DDR1;PDGFRB;IL1A;CCL24;BMP2;WNK2;CCL4;CCL3;CCL2;TREM2;MERTK;FGF10 |
| eosinophil chemotaxis (GO:0048245) | 4/25 | 0.0005209 | 0.034253408 | CCL24;CCL4;CCL3;CCL2 |
| eosinophil migration (GO:0072677) | 4/25 | 0.0005209 | 0.034253408 | CCL24;CCL4;CCL3;CCL2 |

Table S7. GO terms enriched for early events. Cont.
| Term | Overlap | P-value | Adjusted P-value | Genes |
| --- | --- | --- | --- | --- |
| insulin-like growth factor receptor signaling pathway (GO:0048009) | 3/11 | 0.00052917 | 0.034253408 | GHR;IRS1;IGF1 |
| macrophage chemotaxis (GO:0048246) | 3/11 | 0.00052917 | 0.034253408 | EDNRB;CCL3;CCL2 |
| macrophage migration (GO:1905517) | 3/11 | 0.00052917 | 0.034253408 | EDNRB;CCL3;CCL2 |
| positive regulation of MAPK cascade (GO:0043410) | 13/290 | 0.0005437 | 0.034253408 | DDR1;PDGFRB;CCL24;TREM2;IGF1;MERTK;IL1A;BMP2;CCL4;CCL3;CCL2;ELANE;FGF10 |
| positive regulation of transport (GO:0051050) | 5/45 | 0.00059169 | 0.035482639 | C3;DAB2;NR4A3;SCIN;CHRM1 |
| positive regulation of glucose transport (GO:0010828) | 4/26 | 0.00060827 | 0.035482639 | C3;NR4A3;IRS1;IGF1 |
| response to interferon-gamma (GO:0034341) | 6/69 | 0.00064103 | 0.03605796 | CCL24;CCL4;CCL3;CCL2;ACOD1;AQP4 |
| response to tumor necrosis factor (GO:0034612) | 7/98 | 0.00075794 | 0.040696323 | CCL24;DCSTAMP;CCL4;CCL3;CCL2;HAS2;ACOD1 |
| cytokine-mediated signaling pathway (GO:0019221) | 21/634 | 0.00077517 | 0.040696323 | CCL24;EGR1;IL1RN;CSF3;IL1R1;IRS1;BGN;ISG15;PTGS2;CXCL3;CXCL2;IL1A;CXCL12;IL7;IL23A;CCL4;CHAD;CCL3;CCL2;PRTN3;IL9R |
| positive regulation of calcium-mediated signaling (GO:0050850) | 4/29 | 0.00093225 | 0.047364148 | CCL4;CCL3;TREM2;IGF1 |

**Table S8.**
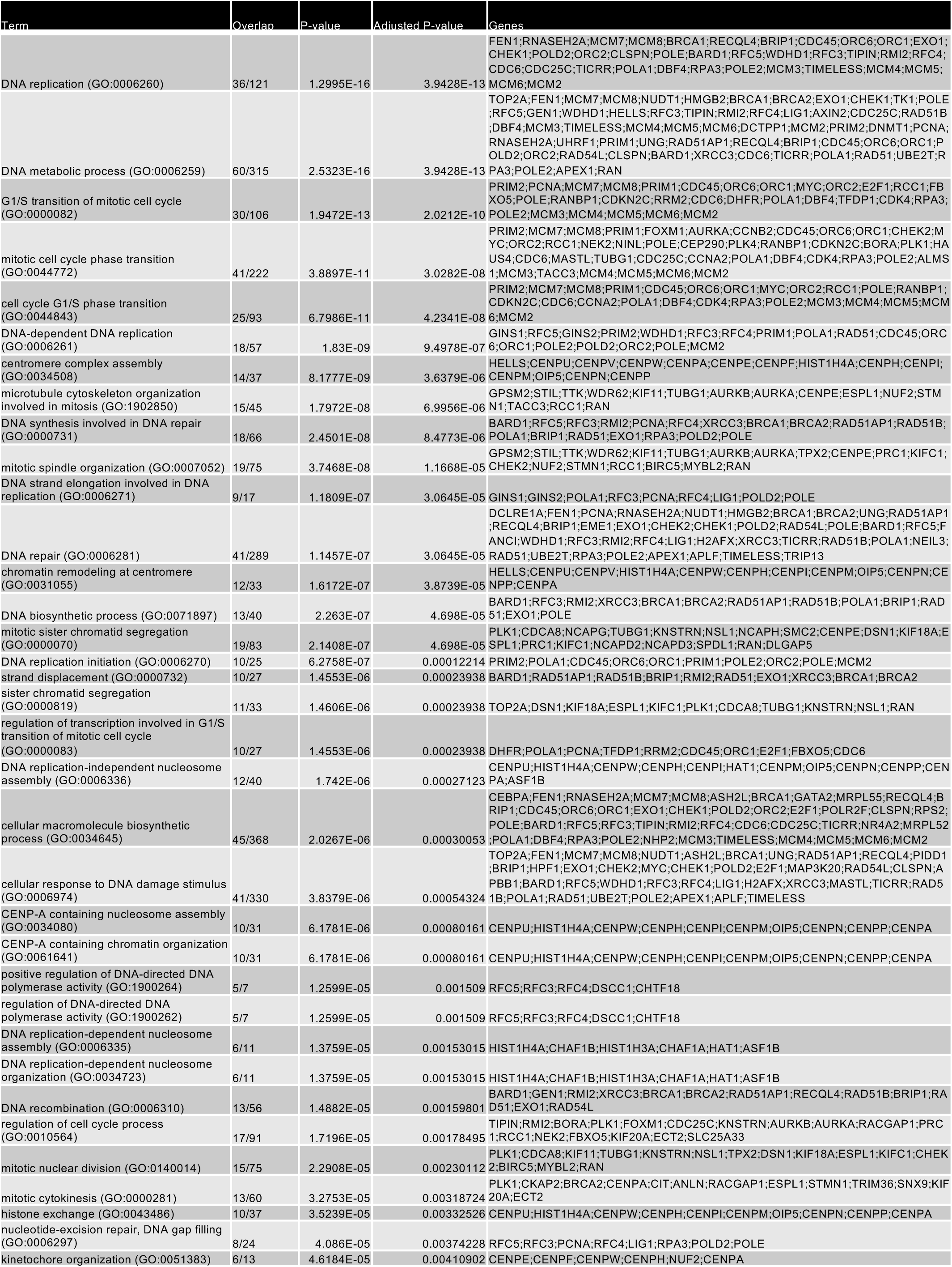
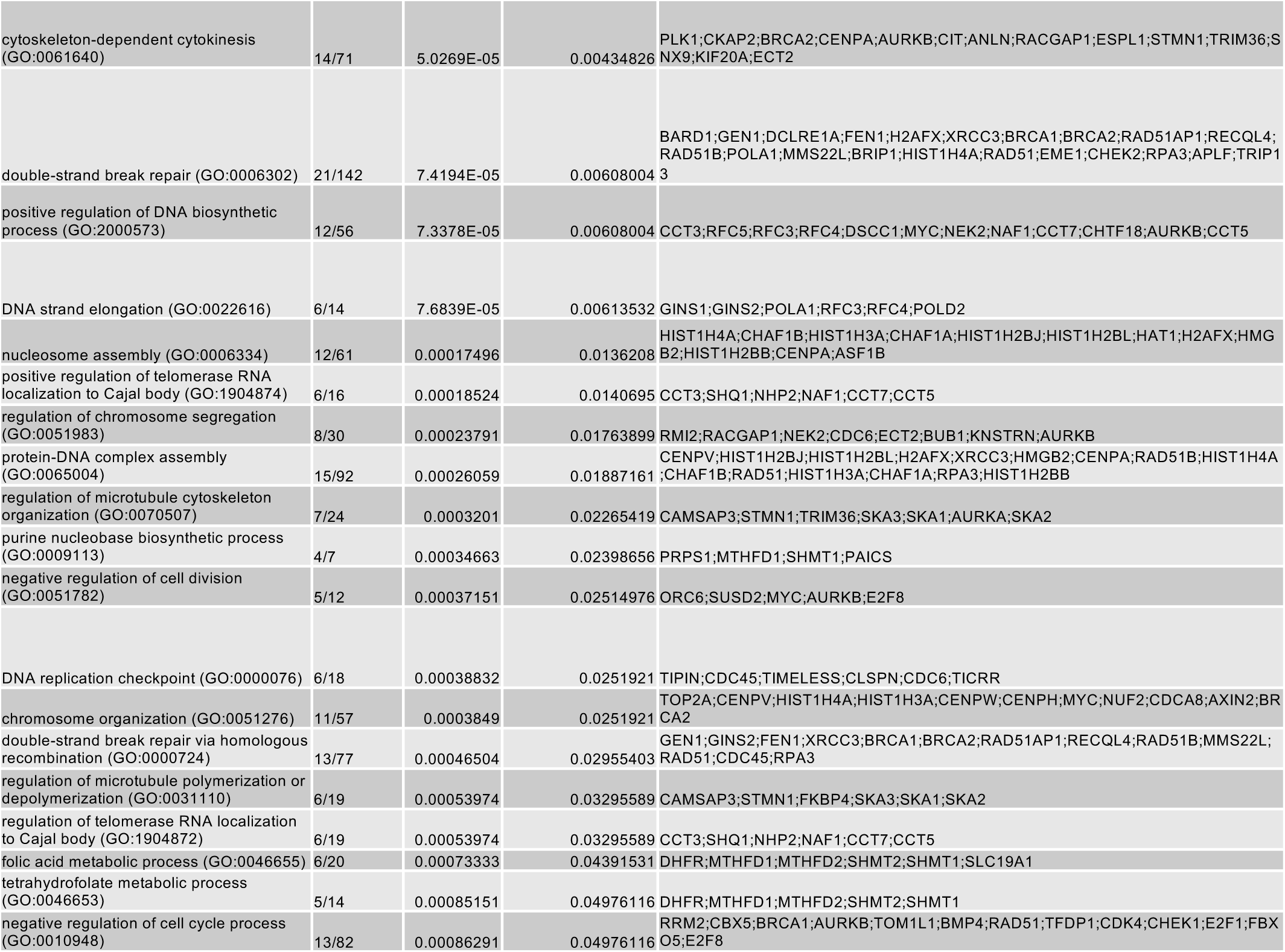
GO terms enriched for transition events.

**Table S9.**
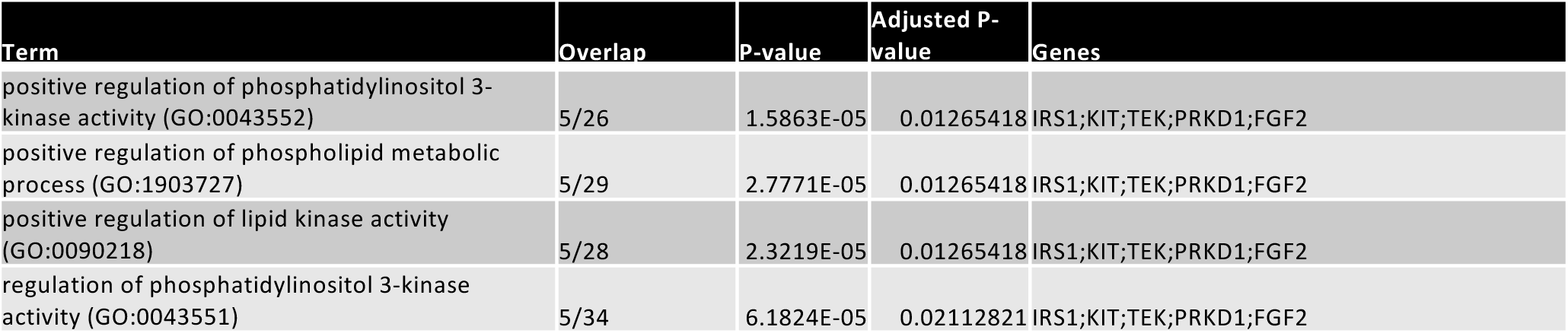
GO terms enriched for persistent events.

| Term | Overlap | P-value | Adjusted P-value | Genes |
| --- | --- | --- | --- | --- |
| positive regulation of phosphatidylinositol 3-kinase activity (GO:0043552) | 5/26 | 1.5863E-05 | 0.01265418 | IRS1;KIT;TEK;PRKD1;FGF2 |
| positive regulation of phospholipid metabolic process (GO:1903727) | 5/29 | 2.7771E-05 | 0.01265418 | IRS1;KIT;TEK;PRKD1;FGF2 |
| positive regulation of lipid kinase activity (GO:0090218) | 5/28 | 2.3219E-05 | 0.01265418 | IRS1;KIT;TEK;PRKD1;FGF2 |
| regulation of phosphatidylinositol 3-kinase activity (GO:0043551) | 5/34 | 6.1824E-05 | 0.02112821 | IRS1;KIT;TEK;PRKD1;FGF2 |

**Table S10.**
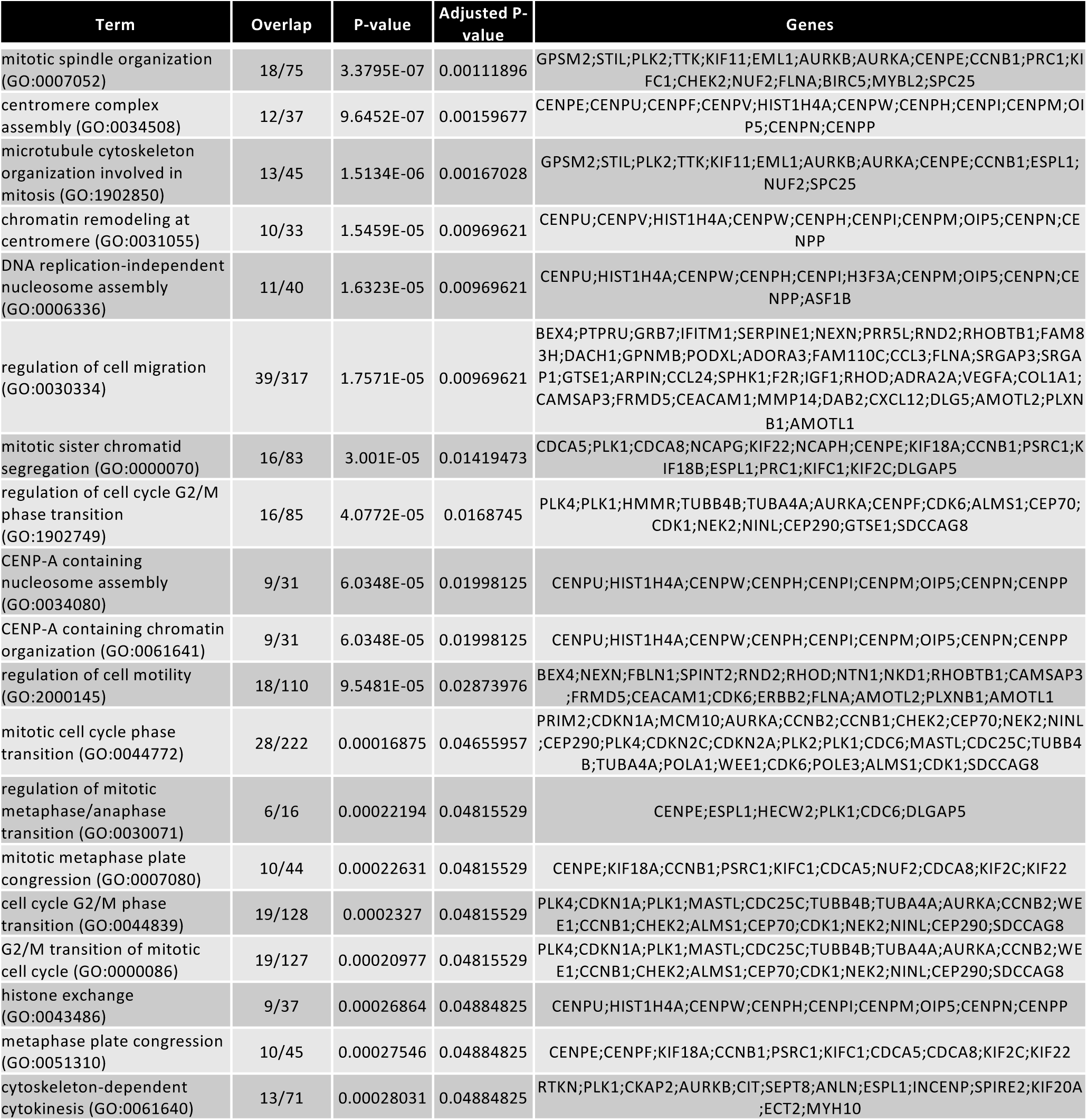
GO terms enriched for leukemia eigenDEG.

